# Contemporary ecological heterogeneity shapes candidate adaptive genomic variation across the native range of an invasive herbivore

**DOI:** 10.64898/2026.09.10.749826

**Authors:** Daniel Poveda-Martínez, Mamionah N. J. Parany, Nicolas A. Salinas, Nicolas N. Moreyra, Esteban Hasson, Laura Varone

## Abstract

Ecological heterogeneity within a species’ native range can shape genomic variation available for subsequent range expansion and biological invasion. The cactus moth, *Cactoblastis cactorum*, is a South American oligophagous herbivore on *Opuntia* that has become an invasive pest outside its native range. In Argentina, populations span wide climatic and geographic gradients and exploit both native *Opuntia* species and the introduced crop *O. ficus-indica*. Using ddRADseq data from 136 individuals across 28 populations, we combined genome-wide differentiation scans (*XtX*), a host-use contrast, and genotype-environment association (GEA) analyses to identify candidate genomic signatures associated with contemporary ecological variation. *XtX* analyses detected 12 candidate regions, whereas the host-use contrast identified 17 regions differentiating populations sampled from *O. ficus-indica* and native *Opuntia*, several of which persisted in a geographically restricted sensitivity analysis. GEA analyses identified 167 covariate-specific loci, including candidate genes associated with detoxification and metabolism (*CYP6B2*), oxygen-sensing pathways (*Egln1*), and circadian regulation (*TIMELESS*). Convergence among analyses was limited but stronger than expected by chance. Temperature-associated loci in *fax* and *Bag6* occurred within an *XtX* candidate region, whereas *Hspg2* was independently recovered in host-use and environmental analyses and retained after geographic restriction. Most candidate variants occurred in non-coding genomic contexts, protein-altering variants were uncommon, and candidate genes spanned diverse functions, consistent with a potentially regulatory and polygenic architecture of adaptation. Overall, our results reveal a heterogeneous genomic landscape associated with climatic variation and introduced-host use, with recurrent loci providing the strongest evidence for localized adaptive differentiation across the native range of *C. cactorum*.

## Introduction

Understanding why some species successfully establish and spread beyond their native ranges remains a central question in evolutionary ecology. Invasion success depends not only on ecological conditions encountered after introduction, but also on the demographic and evolutionary history of source populations before colonization (Sakai et al., 2001; Lee, 2002; Estoup et al., 2016). Introduced populations may establish despite demographic bottlenecks when they retain sufficient standing genetic variation, experience admixture through multiple introductions, or respond rapidly to selection in novel environments, highlighting the importance of the genetic variation carried from the native range (Dlugosch and Parker, 2008; Dlugosch et al., 2015; Estoup et al., 2016). Comparatively, less attention has been given to ecological change occurring within the native range before introduction. Human-mediated habitat modification can alter resource availability, biotic interactions, and abiotic conditions, potentially changing both demographic trajectories and the selective environments experienced by native populations (Estoup et al., 2026). This idea is central to the anthropogenically induced adaptation to invade hypothesis (Hufbauer et al., 2012; Estoup et al., 2026), which proposes that adaptation to human-modified environments within the native range may precondition populations for establishment in similarly altered environments elsewhere. Testing this hypothesis ultimately requires comparisons between native and introduced populations, but its broader premise highlights an important and still underexplored question, how does anthropogenic ecological change reshape genomic variation within the native range itself? Therefore, characterizing this variation is a necessary first step toward understanding the adaptive genomic variation present within native populations before any subsequent range expansion.

This possibility is especially relevant for herbivorous insects, in which host-plant use is a major source of ecological differentiation and can generate strong selection on traits involved in host recognition, feeding, detoxification, development, and reproductive performance (Matsubayashi et al., 2010; Simon et al., 2015; Vertacnik and Linnen, 2017). The incorporation of a novel host into the native range can therefore expose populations to a new combination of chemical, nutritional, phenological, and structural conditions, creating an additional selective environment within an otherwise familiar geographic landscape (Carroll and Boyd, 1992; Carroll et al., 1997). Genomic studies of phytophagous insects show that the genetic basis of host-associated adaptation can vary substantially among traits and systems, ranging from a few loci of relatively large effect to more polygenic architectures involving multiple loci, while host-associated genomic differentiation may arise despite ongoing gene flow (Matsubayashi et al., 2010; Simon et al., 2015; Vertacnik and Linnen, 2017). This makes recently introduced host plants particularly useful for studying how anthropogenic ecological change can leave genomic signatures within native populations. If host use affects survival, development, or reproduction, it may alter allele frequencies at loci involved in host-related traits and influence local demographic dynamics; changes in host availability and distribution may, in turn, affect opportunities for gene flow among populations (Awmack and Leather, 2002; Matsubayashi et al., 2010). Thus, novel host association within the native range provides a direct route by which human-mediated environmental change may reshape both selection and the spatial distribution of standing genetic variation before any later range expansion.

The South American cactus moth, *Cactoblastis cactorum* Berg (Lepidoptera: Pyralidae), provides a useful system in which to examine how anthropogenic changes in host availability interact with environmental heterogeneity within a native range. *Cactoblastis cactorum* is an oligophagous herbivore whose entire life cycle is associated with cacti of the genus *Opuntia* (Cactaceae: Opuntioideae), which provide both feeding and reproductive resources (McFadyen, 1985; Varone et al., 2014). Across southern South America, the species occupies environmentally heterogeneous landscapes including arid and semi-arid regions, the more humid Pampas, and the cold, arid environments of the northern Patagonian steppe. In Argentina, it exploits several native *Opuntia* species, including *O. anacantha*, *O. bonaerensis*, *O. elata*, *O. megapotamica*, *O. penicilligera*, *O. quimilo*, and *O. rioplatense*, together with the cultivated prickly pear *O. ficus-indica* (McFadyen, 1985; Varone et al., 2014). The incorporation of *O. ficus-indica* into this host landscape represents a particularly important anthropogenic change. *Opuntia ficus-indica* originated in Mexico and has a long history of domestication and cultivation before becoming one of the most widely distributed cactus crops worldwide (Kiesling, 1988; Griffith, 2004; Inglese et al., 2017; Montenegro et al., 2024). Its establishment in South America introduced a widespread and abundant host resource absent during much of the earlier evolutionary history of *C. cactorum*. The moth has subsequently become an important pest of cultivated *O. ficus-indica* within its native range, where larval feeding can reduce plant growth and fruit production and, under severe infestation, cause plant mortality and substantial damage to plantations (Fuentes Corona et al., 2025; Varone et al., 2012).

Extensive field surveys across Argentina showed that realized host use by *C. cactorum* largely tracks local *Opuntia* availability, with both native and introduced hosts exploited across the native range (Varone et al., 2014). *Opuntia ficus-indica* was among the most geographically widespread hosts surveyed, whereas several native *Opuntia* species had more restricted distributions. Laboratory assays, nevertheless, detected differences in oviposition preference, including preference for *O. ficus-indica* under some conditions (Varone et al., 2014), and laboratory performance studies showed that *C. cactorum* generally developed more rapidly on *O. ficus-indica* than on several native *Opuntia* hosts (Varone et al., 2012).

Previous studies indicate that genetic variation within the native range of *C. cactorum* has been shaped by a combination of geography, demographic history, and environmental change. Mitochondrial analyses revealed high haplotype diversity and pronounced geographic structure across the native range (Marsico et al., 2011), while phenotypic studies further documented regional ecotypic variation, including differences in larval coloration and host-related performance among native populations (Brooks et al., 2012). More recently, landscape-genomic analyses showed that geography, Quaternary climatic oscillations, historical shifts in the distribution of *Opuntia* hosts, and contemporary environmental variation contributed to genomic structure across the native range (Andraca-Gómez et al., 2024; Poveda-Martínez et al., 2023). Notably, these genomic analyses did not identify contemporary host use as a major determinant of broad population structure. Thus, although the native genomic landscape retains a strong historical and spatial signature, it remains unclear whether more recent ecological pressures, including use of the anthropogenically introduced *O. ficus-indica* and present-day climatic gradients, have generated localized genomic differentiation superimposed on this background.

This question is particularly relevant given the subsequent invasion history of *C. cactorum*. Individuals originating from Argentina were intentionally introduced into Australia during the 1920s for biological control of invasive prickly pears, and the species was subsequently established through additional deliberate and accidental introductions in regions including South Africa, the Caribbean, and North America (Zimmermann et al., 2000; Marsico et al., 2011). Although its impact in Australia became a well-known example of successful biological control, *C. cactorum* is now also regarded as a major threat to native and cultivated *Opuntia* in regions where it has spread beyond intended control programmes, particularly in the Caribbean and North America (Hight et al., 2005; Bloem et al., 2005; Zimmermann et al., 2000; Varone et al., 2020). Genetic studies initially helped reconstruct these introduction histories and identify South American source regions (Marsico et al., 2011), while subsequent work increasingly resolved genomic variation within the native range (Andraca-Gómez et al., 2024; Poveda-Martínez et al., 2023). Thus, while the historical and spatial organization of native populations is increasingly well resolved, the extent to which contemporary ecological heterogeneity has driven local adaptation in *C. cactorum* remains largely unknown. From the perspective of the anthropogenically induced adaptation to invade hypothesis, characterizing whether human-mediated ecological change has already shaped genomic variation within source populations represents an important step before testing whether such variation subsequently contributes to invasion success.

Here, we investigate genomic signatures associated with contemporary ecological heterogeneity across the native range of *C. cactorum* in Argentina. Using genome-wide SNP data from populations sampled across climatic, geographic, and host-use gradients, we combine complementary differentiation and genotype-environment association approaches to test whether (i) particular genomic regions show elevated differentiation beyond the genome-wide background; (ii) populations predominantly associated with *O. ficus-indica* show localized genomic differentiation relative to populations associated with native opuntias, and whether these signals persist after geographic restriction; and (iii) allele frequencies at particular loci are associated with climatic and/or spatial gradients. We hypothesize that broad genomic structure mainly reflects historical and geographic differentiation, whereas contemporary ecological responses are concentrated in a more restricted subset of loci associated with host use and environmental variation. Because the different analyses capture complementary dimensions of genomic variation, we further expect that recurrent signals across approaches provide particularly strong candidates for local adaptation. By resolving these native-range patterns, our study provides a foundation for future comparisons with introduced populations and for testing whether standing ecological variation in South America contributed to the evolutionary potential of *C. cactorum* during subsequent range expansion.

## Material and Methods

### Dataset and study area

We analysed ddRADseq genomic data from 136 *C. cactorum* individuals sampled across the species’ native range in Argentina (Fig. 1A; Table S1). These individuals correspond to a subset of the material originally generated by Poveda-Martínez et al. (2023). Briefly, between 2018 and 2019, individuals were collected from 28 populations distributed across 13 Argentine provinces and spanning three major biogeographic regions: the Monte, Chacoan, and Pampean. Sampling covered a broad geographic extent, from Jujuy Province in the northwest (-24.39357, -65.27129) and Chaco Province in the northeast to Río Negro Province, northern Patagonia, in the south (-39.35746, -65.71550), and from La Pampa Province in the west (-36.23290, -66.93740) to Entre Ríos Province in the east (-30.65441, -58.00371). The sampling area encompasses marked environmental heterogeneity, including broad climatic, topographic, and ecological gradients. Arid and semi-arid environments characterize much of the northern and western portions of the sampled range, whereas grassland and steppe environments are more characteristic in eastern and southern localities, respectively.

**Figure 1.**
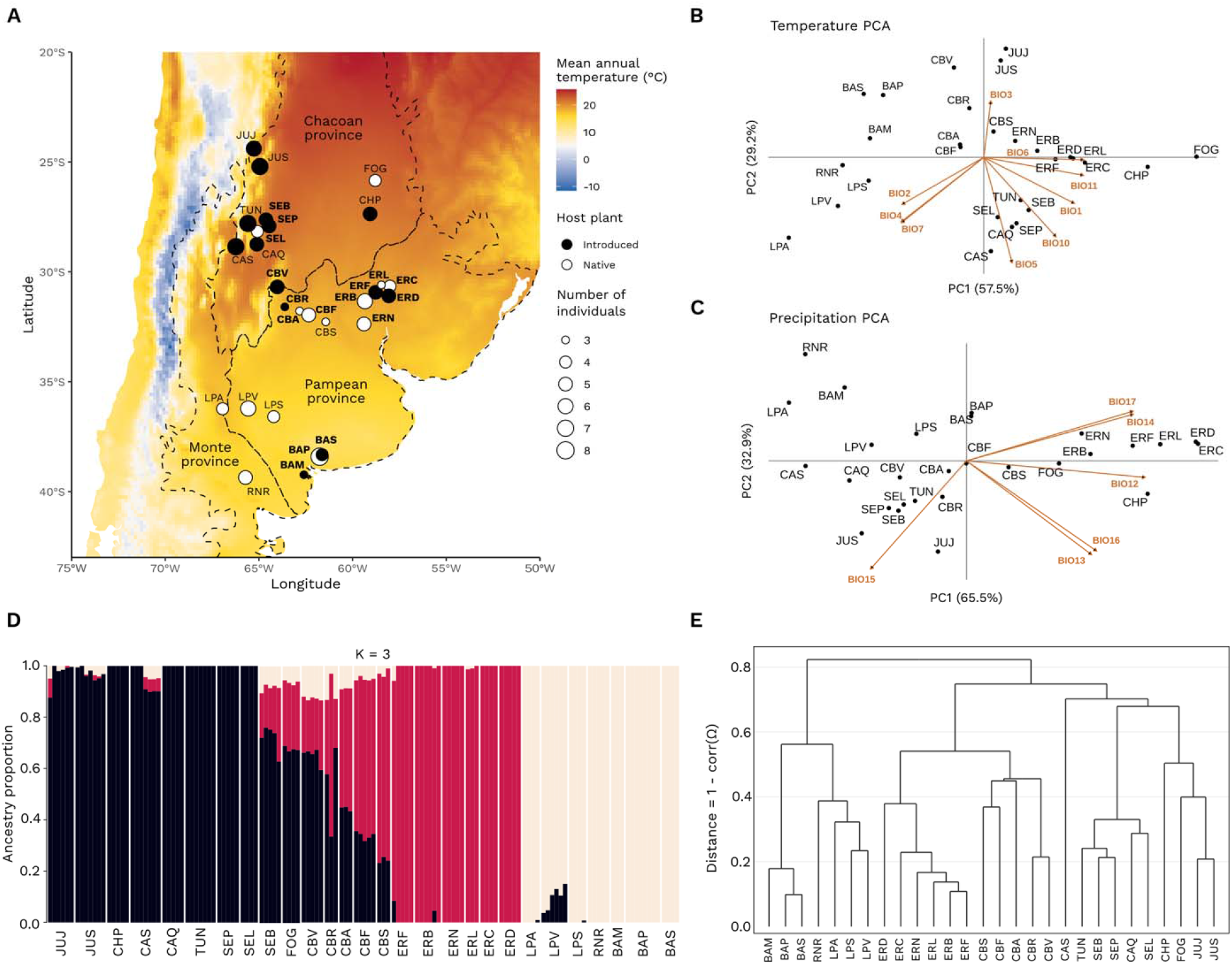
Sampling design, climatic variation, population structure, and genomic covariance across the native range of *Cactoblastis cactorum* in Argentina. **(A)** Geographic distribution of the 28 sampled populations across the native range, showing host-use category, mean annual temperature (BIO1), and the main biogeographic regions represented in the study following Morrone (2014) regionalization. Populations were classified according to whether individuals were sampled predominantly from the introduced host *Opuntia ficus-indica* or from native *Opuntia* species. The 16 populations retained for the geographically restricted sensitivity analysis of the *C_2_* host-use contrast are indicated in bold. **(B)** Principal component analysis (PCA) of temperature-related bioclimatic variables (BIO1-BIO7, BIO10, and BIO11). TemperaturePC1 explained 57.5% of the total variation and represented a composite thermal gradient. **(C)** PCA of precipitation-related bioclimatic variables (BIO12-BIO17). PrecipitationPC1 explained 65.5% of the total variation. In panels B and C, points represent sampling populations and arrows indicate the direction and relative contribution of individual bioclimatic variables. TemperaturePC1 and PrecipitationPC1 were retained as synthetic climatic covariates for downstream genotype-environment association analyses. **(D)** ADMIXTURE ancestry proportions at (*K*= 3), highlighting the three major geographic lineages previously described across the native range of *C. cactorum*. The best-supported solution based on cross-validation was (*K*= 6) and is shown together with the full range of (*K*) values in Fig. S5. Each vertical bar represents one individual, partitioned according to its estimated ancestry in the corresponding genetic clusters. (**E**) Dendrogram derived from the BayPass population covariance matrix (Ω), summarizing covariance among populations and recapitulating the major genetic structure of the dataset. This covariance structure was incorporated into BayPass analyses to account for shared demographic history and reduce confounding between population structure and signals of selection.

Larvae were collected directly from host plants, including seven native *Opuntia*: *O. anacantha, O. bonaerensis, O. elata, O. megapotamica, O. penicilligera, O. rioplatense*, and *O. quimilo* and the exotic *O. ficus-indica*. To minimize biases associated with close relatedness and uneven sampling, only one individual per sampled plant was retained for genetic analyses. After read processing, mapping, and quality control, we retained 136 individuals from 28 populations, with 3-8 individuals per population, for downstream population genomic analyses. Genomic libraries were prepared following the double-digest restriction-site associated DNA sequencing protocol described by Peterson et al. (2012), using the restriction enzymes *NspI* and *MboI*. Individuals were sequenced in paired-end 125 bp mode on an Illumina HiSeq2500 platform. SRA accessions, sampling coordinates, host plant information, and additional geographic and ecological metadata are provided in Table S1.

### ddRADseq processing, mapping and variant calling

Raw paired-end ddRADseq reads were first assessed with FastQC, and quality-control reports were summarized with MultiQC (Ewels et al., 2016). Reads were then processed with process_radtags from Stacks v2.68 (Rochette et al., 2019) using a paired-end configuration. Reads with uncalled bases or low-quality scores were discarded, and Illumina adapter contamination was also removed. Processed reads were aligned to the *C. cactorum* reference genome with bwa-mem2 v2.2.1 (Vasimuddin et al., 2019). Variant calling was performed using the reference-based workflow implemented in Stacks. BAM files were processed with gstacks to identify RAD loci and call SNPs and genotypes across individuals, using a minimum mapping-quality threshold of 10. Population-level filtering was then performed with the *Populations* module. Loci were retained if present in at least two populations and genotyped in at least 90% of individuals within those populations (-*p 2 -r 0.90*). The resulting VCF was further processed using VCFtools v0.1.16 (Danecek et al., 2011) and BCFtools v1.19 (Danecek et al., 2021).

Missing data were quantified at both the individual and SNP levels using *--missing-indv* and -- *missing-site*, respectively. Subsequent filtering retained only biallelic SNPs, excluded loci with more than 20% missing genotypes (--*max-missing 0.80*), and retained variants with a minor allele frequency ≥ 0.05. The number of SNPs retained after each filtering step is reported in Table S2. The all-SNP dataset comprised 132,283 biallelic SNPs genotyped in at least 80% of individuals. To obtain a second dataset with reduced linkage disequilibrium for analysis requiring approximately independent markers, the all-SNP dataset was pruned using Plink2 (Chang et al., 2015) with a sliding window of 50 variants, a step size of 10 variants, and an r² threshold of 0.20 (--*indep-pairwise 50 10 0.2*). This procedure retained 48,343 linkage-independent SNPs, hereafter referred to as the unlinked-SNP dataset. This dataset was used for population-structure analysis that required reduced linkage among markers, whereas the all-SNP dataset was retained for genome-scan analyses to maximize marker density and genomic resolution.

### Environmental and spatial covariates

Environmental and spatial covariates were compiled for each population and prepared for genotype-environment association analyses. To characterize climatic variation across the native range of *C. cactorum*, we extracted 19 bioclimatic variables from WorldClim v2.1 at 2.5-arc-minute resolution, corresponding to approximately 5 km at the equator (Fick and Hijmans, 2017). Following recommendations to avoid known spatial artefacts in some WorldClim variables, we excluded BIO8, BIO9, BIO18, and BIO19 (Oliveira et al., 2020), resulting in 15 climatic predictors. These included annual and seasonal components of temperature (BIO1-BIO7; BIO10-BIO11) and precipitation (BIO12-BIO17) variation (see Table S3).

To reduce dimensionality and collinearity among climatic predictors while retaining biologically interpretable environmental gradients, temperature- and precipitation-related bioclimatic variables were analysed in two separate principal component analyses (PCA). The first principal component of temperature variables (TemperaturePC1) explained 57.5% of total thermal variation among populations (Fig. 1B), with major contributions of minimum temperature of the coldest month (BIO6), mean temperature of the coldest quarter (BIO11), annual mean temperature (BIO1), temperature seasonality (BIO4), annual temperature range (BIO7), and mean diurnal range (BIO2). The first principal component of precipitation variables (PrecipitationPC1) accounted for 65.5% of total variation in precipitation and was primarily associated with annual precipitation (BIO12), precipitation in the driest month (BIO14), precipitation in the driest quarter (BIO17), precipitation in the wettest quarter (BIO16), and precipitation in the wettest month (BIO13) (Fig. 1C; Fig. S1; Table S3). TemperaturePC1 and PrecipitationPC1 were retained as synthetic climatic covariates summarizing multivariate thermal and precipitation gradients. We additionally included an aridity index because water availability and evaporative demand are ecologically relevant to the arid and semi-arid environments inhabited by *Opuntia* hosts and, consequently, the cactus moth. Annual Aridity Index is the ratio of annual precipitation to reference potential evapotranspiration, with lower values corresponding to more arid conditions. Values of this index were obtained from the Global Aridity Index and Potential Evapotranspiration Database v3 at 30-arc-second resolution (Zomer et al., 2022). We also included elevation as an additional environmental predictor because altitudinal gradients can influence local thermal conditions and covary with moisture, season length, vegetation, and other ecological factors (Körner, 2007). Elevation values were extracted at 30-arc-second resolution using the geodata R package (Hijmans et al., 2026). Finally, spatial gradients were represented by latitude and longitude, which were included as continuous covariates to capture broad geographic structure across the sampled range.

To characterize covariate structure and potential collinearity, we computed pairwise Pearson correlations among the final quantitative environmental and spatial predictors, TemperaturePC1, PrecipitationPC1, aridity index, elevation, latitude, and longitude (Fig. S2). Several predictors were strongly correlated, particularly PrecipitationPC1 with aridity index and longitude. However, these correlations do not introduce multicollinearity within the genotype-environment association models because each covariate was tested independently in a separate standard covariate model, with only one environmental or spatial predictor included at a time. The correlation analysis was therefore used primarily to characterize redundancy among environmental gradients and to guide biological interpretation of overlapping genomic signals across covariates. The final set of environmental and spatial variables used in genotype-environment association analyses is summarized in Table S4.

### Population structure

Population structure was investigated using the unlinked-SNP dataset with ADMIXTURE v1.3.0 (Alexander et al., 2009), which uses a maximum-likelihood framework to estimate individual ancestry proportions without prior assignment of individuals to populations. ADMIXTURE was run in unsupervised mode for values of *K* ranging from 2 to 8, where *K* represents the assumed number of ancestral genetic components. Model fit across values of *K* was evaluated using the cross-validation procedure implemented in ADMIXTURE, with lower cross-validation error indicating better predictive fit. The value of *K* with the lowest cross-validation error was considered the best-supported representation of population structure, while ancestry patterns across the full range of *K* values were also examined.

### Detecting signatures of selection

#### BayPass framework and genome-wide differentiation

Genome-wide scans of population differentiation, host-associated allele frequency differentiation, and genotype-environment association (GEA) were performed using BayPass v3.1 (Gautier, 2015). BayPass models the genome-wide covariance in allele frequencies among populations through the population covariance matrix Ω, thereby accounting for shared demographic history and population structure when identifying loci showing unusually high differentiation or association with population-level covariates (Gautier, 2015). This framework is particularly appropriate for landscape genomic analyses in structured populations because it reduces the risk of false positives caused by neutral population structure. Analyses were conducted using population-level allele count matrices generated from the all-SNP dataset. BayPass was first run under the core model to estimate the population covariance matrix (Ω) and to compute the *XtX* statistic (Olazcuaga et al., 2020). *XtX* is a Bayesian differentiation statistic that identifies SNPs showing stronger allele frequency differentiation than expected under the genome-wide covariance structure among populations (Gautier, 2015; Olazcuaga et al., 2020). This analysis was used to detect candidate loci showing unusually high differentiation across the native range of *C. cactorum*, after accounting for shared demographic history and population structure.

#### Host-use contrast and geographic sensitivity analysis

To evaluate genomic differentiation associated with host use, populations were classified according to the predominant host category represented among sampled individuals. Fourteen populations were assigned to the native-*Opuntia* group (*O. anacantha, O. bonaerensis, O. elata, O. megapotamica, O. penicilligera, O. rioplatense*, and *O. quimilo*) and 14 to the introduced-host group (O. ficus-indica) (Fig. 1A; Table S4). At five sampling sites, individuals were collected from both native and introduced host categories; however, one category clearly predominated at each site, with only a single sampled plant belonging to the alternative category. These populations were therefore assigned according to the dominant host category at the site. For the BayPass contrast, native-host populations were coded as −1 and *O. ficus-indica*-associated populations as +1. Host-associated allele-frequency differentiation was assessed using the *C_2_* contrast statistic (Olazcuaga et al., 2020) implemented in BayPass. The *C_2_* statistic evaluates differentiation between predefined population groups while accounting for genome-wide covariance in allele frequencies through the population covariance matrix (Ω).

Because host categories are spatially structured across the sampled range, we additionally evaluated the robustness of the host-associated signal using a geographically restricted sensitivity analysis. This analysis included 16 populations from Buenos Aires, Córdoba, Entre Ríos, and Santiago del Estero, the four provinces in which both host categories are present, resulting in a balanced contrast of eight populations sampled from *O. ficus-indica* and eight from native *Opuntia* (Fig. 1A). For this restricted dataset, the population covariance matrix was re-estimated independently before rerunning the *C_2_* contrast. To preserve the SNP ascertainment of the primary dataset while removing sites that became uninformative after population restriction, only loci that remained polymorphic across the 16 retained populations were included in the sensitivity analysis. Concordance with the full-range analysis was evaluated using genome-wide *C_2_*statistics, retention and ranking of primary candidate peaks, physical overlap among candidate regions, and shared peak-associated genes.

#### Genotype-environment association analyses

GEA analyses were performed using the BayPass standard covariate model (Gautier, 2015). Environmental and spatial covariates included TemperaturePC1, PrecipitationPC1, aridity index, elevation, latitude, and longitude. All quantitative covariates were analysed independently, with one predictor per model to avoid fitting correlated predictors within a single multivariate framework. Covariates were standardized within BayPass using the *-scalecov* option, allowing regression coefficients and Bayes Factors (BFs) to be estimated on comparable scales across environmental and spatial predictors (Gautier, 2015). Evidence for association between allele frequencies and each covariate was evaluated using BFs estimated under the standard covariate model.

#### BayPass convergence, candidate-region detection and cross-analysis overlap

Each BayPass analysis (*XtX, C_2_*and GEA) was replicated using three independent Markov chain Monte Carlo (MCMC) chains initiated with different random seeds. To improve numerical precision and assess convergence, each chain consisted of 20 pilot runs of 500 iterations, followed by a burn-in of 10,000 iterations and 2,500 post-burn-in samples recorded every 20 iterations. Convergence and numerical robustness were evaluated using complementary diagnostics. Population covariance matrices (Ω) inferred from independent chains were compared using the Förstner-Moonen distance (FMD; Förstner and Moonen, 2003), as implemented in the BayPass R utility function *fmd.dist*, with low FMD values indicating similar covariance estimates among runs. We additionally compared posterior mean estimates of the Beta-prior parameters governing population allele frequencies and calculated pairwise Spearman rank correlations of SNP-level *XtX* estimates among independent chains. Following convergence, BFs from GEA analyses were extracted independently for each covariate and seed, and summarized as the median across the three MCMC runs to obtain final SNP-level BF estimates. *XtX* and *C_2_* statistics were not combined across seeds; downstream candidate identification was based on a representative converged run selected after confirming stable covariance estimates and strong reproducibility of genome-wide *XtX* values across chains.

To account for linkage and spatial clustering of differentiated SNPs along the reference genome, we applied the local score approach of Fariello et al. (2017) to the BayPass *XtX* and *C_2_* statistics. Local scores were computed independently for each scaffold and restricted to scaffolds >1 Mb. SNP-specific scores were defined as −log10(p) − ξ, where ξ is a penalty parameter controlling the stringency of candidate-region detection, and scores were accumulated along each eligible scaffold to identify genomic regions containing clusters of elevated signals rather than isolated SNP outliers. Low-frequency variants (MAF ≤0.05) were excluded from local-score calculations. We used ξ = 2 and α = 0.01 for both *XtX* and *C_2_*analyses. To provide an independent neutral calibration of the *XtX* and *C_2_* signals, we generated pseudo-observed datasets (PODs) using the function simulate.baypass. We simulated 100,000 SNPs using the population covariance matrix (Ω), Beta-prior parameters, and population sample-size distribution estimated from the empirical data. Simulated SNPs were filtered using the same MAF > 0.05 criterion applied in the local-score analyses, and the 99.9th percentile of the resulting neutral distributions was used as the POD threshold (POD99.9). This corresponded to *XtX* = 57.39 and *C_2_* = 12.08. Local-score regions remained the primary definition of *XtX* and *C_2_* candidates; POD calibration was used to evaluate whether individual SNPs within these regions exceeded expectations under the neutral model.

For GEA analyses, BFs expressed in decibans [BF(dB)] were extracted independently for each environmental covariate from three converged BayPass runs. For each SNP-covariate combination, the final association statistic was defined as the median BF across the three runs. SNPs with a median BF >20 dB were retained as strongly associated variants. Candidate loci were then defined independently for each covariate by grouping retained SNPs located on the same scaffold and separated by no more than 1 kb, thereby consolidating tightly clustered associations likely to reflect the same local genomic signal; within each locus, the SNP with the highest median BF was designated as the peak SNP. Because the aim of the GEA was to identify SNP-level associations with environmental covariates, GEA candidates were defined from replicated BF estimates and local clustering, without applying the local-score procedure used to detect spatially accumulated differentiation signals in the *XtX* and *C_2_*analyses.

To evaluate whether candidate SNPs were shared among analyses more frequently than expected by chance, we used one-sided hypergeometric tests, following a similar approach recently applied to BayPass outlier sets by Nugroho et al. (2026) and based on Fury et al. (2006). For these comparisons, the *XtX* and *C_2_* SNP sets comprised SNPs supported by both criteria, i.e. POD99.9 outliers located within their respective local-score candidate regions, whereas GEA sets comprised SNPs with median BF > 20 dB. We tested overlap between the *XtX* and *C_2_*sets and between each of these sets and the six covariate-specific GEA sets. False discovery rate correction using the Benjamini-Hochberg procedure was applied across the 12 covariate-specific GEA comparisons.

### Candidate genes and functional annotation

Significant *XtX*, *C_2_*, and GEA candidates were annotated against the *C. cactorum* reference genome. Peak coordinates were classified as coding, intronic, gene-proximal (within 5 kb of an annotated gene), or distal intergenic. When a peak did not overlap an annotated gene, the nearest gene within 5 kb was reported. Candidate-gene convergence was evaluated separately from physical overlap among candidate regions. For recurrent candidate genes that remained uncharacterized, predicted protein sequences were queried against the NCBI nr and RefSeq protein databases using BLASTP implemented in NCBI BLAST+ v2.15.0 (Camacho et al., 2009), and putative functions were assigned from the best-supported homologous match; protein features were additionally evaluated using InterProScan v5.73-104.0 (Jones et al., 2014).

Predicted variant consequences were characterized for all unique SNPs in the *XtX*, *C_2_*, and GEA candidate sets using Ensembl Variant Effect Predictor (VEP v111; McLaren et al., 2016). The final annotation comprised 567 unique candidate SNP positions. When multiple consequences were assigned to the same SNP, a single primary consequence was retained using the following hierarchy: protein-altering, UTR, synonymous, non-coding exon/transcript, intronic/splice-region, upstream/downstream, and intergenic. Gene Ontology enrichment was performed with topGO (Alexa and Rahnenführer, 2026) for the *XtX*, *C_2_*, pooled GEA, covariate-specific GEA, and combined candidate-gene sets. Biological Process, Molecular Function, and Cellular Component ontologies were tested using Fisher’s exact test with the *weight01* algorithm. The background comprised genes overlapping or located within 5 kb of SNPs screened in the all-SNP BayPass dataset. Benjamini-Hochberg correction was applied within each analysis and ontology, with FDR < 0.05 considered significant.

## Results

After quality filtering, 1.85 ± 0.67 million paired-end read pairs per individual were retained, representing 99.44 ± 0.41% of the raw data. Alignment to the *C. cactorum* reference genome yielded a mean mapping rate of 98.28 ± 2.63% (Fig. S3). On average, 1.49 ± 0.52 million read pairs were properly paired, corresponding to 80.37 ± 3.04% of retained read pairs. Genome-wide sequencing depth averaged 1.31 ± 0.48×, with 13.94 ± 2.45% of the reference genome covered by at least one read (Fig. S4). Variant calling identified 808,111 SNPs. After retaining biallelic variants genotyped in at least 80% of individuals and with a minor allele frequency ≥ 0.05, the all-SNP dataset comprised 132,283 SNPs (Table S2). Linkage-disequilibrium pruning retained 48,343 independent SNPs for analyses requiring reduced linkage among markers. Individual missingness remained low after filtering, averaging 8.92 ± 5.64% in the all-SNP dataset and 8.27 ± 5.54% in the unlinked-SNP dataset. The latter was used for analyses sensitive to linkage among markers, whereas the all-SNP dataset was used to detect genomic signatures associated with selection.

### Population structure, BayPass convergence and covariance

ADMIXTURE identified *K*=6 as the best-supported clustering solution based on the lowest cross-validation error (Fig. S5). This solution revealed finer-scale ancestry structure, with six spatially organized components corresponding to northern, central, western, southeastern, southwestern, and eastern groups. At *K*=3, the analysis recovered the three major geographically structured lineages previously described: central-northern-western, eastern, and southern lineages (Fig. 1D). Admixture was particularly evident among northern and central populations, and remained visible across several *K* values.

Because subsequent BayPass analyses explicitly accounted for shared population history through the covariance matrix Ω, we first evaluated the stability of this inferred structure across independent MCMC runs. Ω estimates were highly consistent among seeds, with low pairwise Förstner-Moonen distances (FMD = 0.076-0.086; Table S5; Fig. S6). SNP-level *XtX* estimates were similarly reproducible, with pairwise Spearman correlations of ρ=0.974 across runs, and posterior estimates of the Beta-prior parameters were also closely concordant (Table S5; Fig. S6). These diagnostics indicated stable and robust BayPass inference.

The inferred Ω matrix itself recovered a pronounced geographic covariance structure that was concordant with the ancestry patterns detected with ADMIXTURE (Fig. 1E). Hierarchical clustering and Ω-based ordination distinguished the southern assemblage comprising populations from Buenos Aires, La Pampa, and Río Negro, an eastern assemblage including populations from Entre Ríos, Santa Fe and part of Córdoba, and a central-northern assemblage comprising populations from Catamarca, Santiago del Estero, Salta, Formosa, and Jujuy. Thus, the covariance structure captured the same major geographic relationships identified by the independent population-structure analysis while also providing the demographic background used to control for shared population history in subsequent BayPass selection and association analyses.

### Genomic signatures of divergent selection and host-use differentiation

The *XtX* genome scan identified 12 candidate regions across seven scaffolds, spanning 597.70 kb in total (Fig. 2A; Table S6). The length of the regions ranged from 2.64 to 177.56 kb and contained 8-47 SNPs. POD calibration strongly supported the local-score results, all 12 regions contained at least one SNP exceeding the POD99.9 threshold, and the original BayPass statistic peak exceeded this threshold in every region; the local-score peak itself exceeded POD99.9 in 11 of the 12 regions. In total, the retained regions contained 57 POD99.9-supported SNPs. All 12 candidate peaks were located within or near annotated genes: six were intronic, three were gene-proximal (within 5 kb of an annotated gene), and three fell within coding sequence. The strongest local-score signal occurred on scaffold ptg000103 within a gene annotated as Techylectin-5B. Other prominent candidates involved *lmx1b.1*, *ARL5B*, *Bag6*, *Phgdh*, *KCNT1*, *Neurl4*, *mep-1*, *TBC1D13*, and an uncharacterized gene, *Ccac_g11528*, which showed strong sequence similarity to lepidopteran *Masc* gene that encodes *Masculinizer* proteins (best BLASTP hit: 56.9% identity over 84.1% of the query, *E* = 2.17 x 10^-87^) and was therefore annotated as putative *Masc* homolog. The broader *XtX* region on scaffold ptg000084 also encompassed *fax*. Candidate genes were associated with functions related to signalling, metabolism, ion transport, neural and developmental processes, immune function, and sex determination.

**Figure 2.**
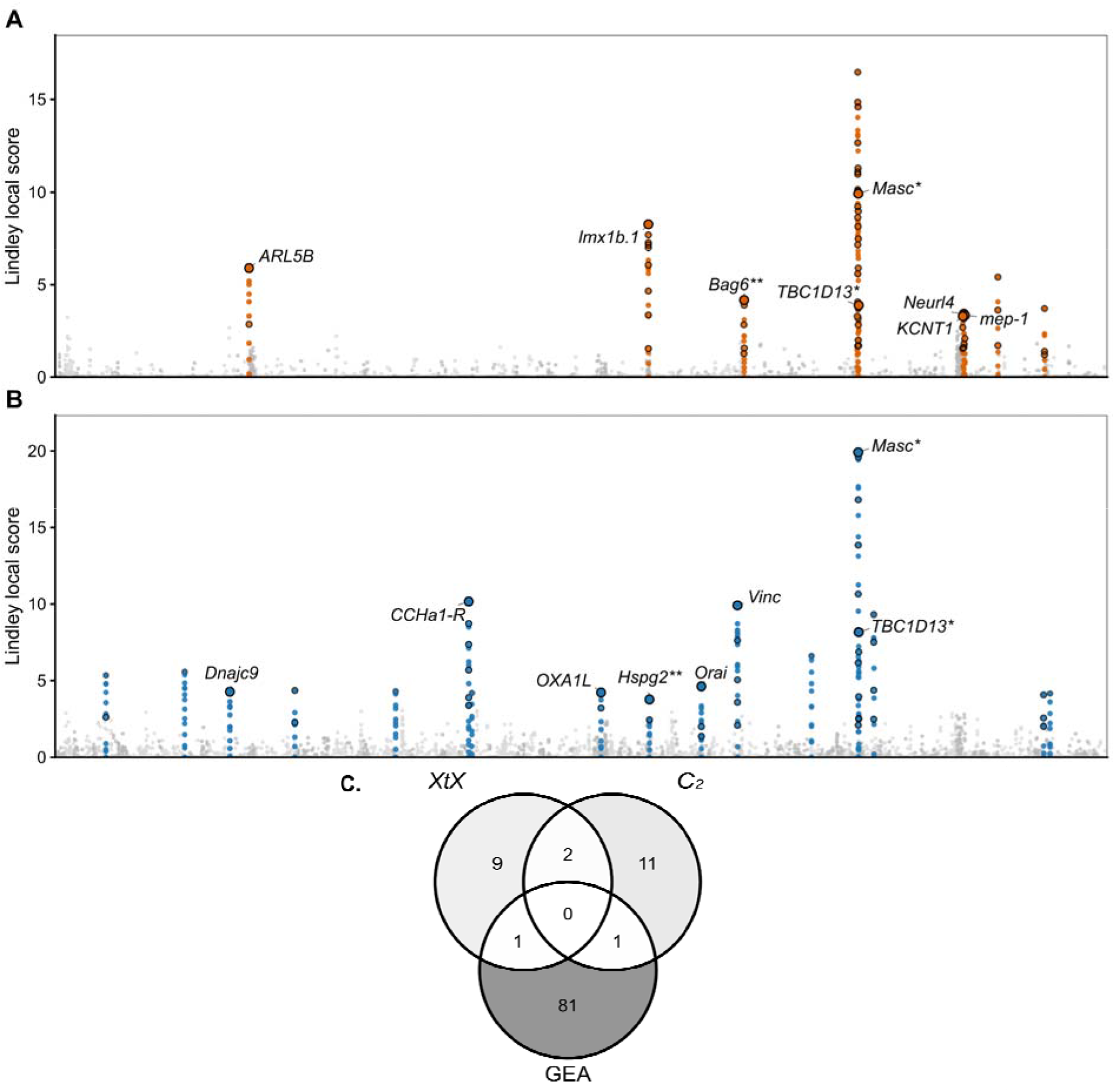
Genome-wide differentiation, host-associated differentiation, and overlap among candidate genes. (**A**) Genome-wide differentiation identified using BayPass *XtX* statistics and the local-score approach. SNP-wise local scores are plotted across concatenated genomic scaffolds >1 Mb, with candidate regions highlighted in orange and selected annotated genes labelled. (**B**) Host-associated differentiation identified using the BayPass *C_2_* contrast for host use, with candidate regions highlighted in blue and selected annotated genes labelled. In both panels, background SNPs are shown in grey tones, and open black circles indicate SNPs within retained local-score candidate regions that also exceeded the POD99.9 threshold. The x-axis represents genomic position across concatenated scaffolds. A single asterisk (*) identifies genes shared between the *XtX* and *C_2_* candidate sets, whereas two asterisks (**) identify genes also recovered in GEA analyses. Thus, *Masc* and *TBC1D13* were shared between *XtX* and *C_2_*, *Bag6* was shared between *XtX* and GEA, and *Hspg2* was shared between *C_2_* and GEA. (**C**) Venn diagram showing overlap among unique genes associated with *XtX*, *C_2_*, and GEA candidates. The three analyses identified 12, 14, and 83 unique candidate genes, respectively. Two genes were shared between *XtX* and *C_2_*, one between *XtX* and GEA, and one between *C_2_* and GEA, with no gene shared among all three analyses. Gene-level overlap does not necessarily imply physical overlap of the corresponding candidate genomic intervals.

The *C_2_* host-use contrast identified 17 candidate regions across 15 scaffolds, spanning 603.77 kb in total (Fig. 2B; Table S7). Retained regions ranged from 60 bp to 138.29 kb. POD calibration supported a substantial subset of these signals. Eleven out of the 17 regions contained at least one SNP exceeding POD99.9, and the original BayPass statistic peak exceeded this threshold in the same 11 regions, whereas the local-score peak itself exceeded POD99.9 in seven regions. Collectively, the retained regions contained 38 unique POD99.9-supported SNPs. The strongest local-score signal occurred on scaffold ptg000103 near the putative *Masc* homolog, with additional prominent candidates including *CCHa1-R*, *Vinc*, *Hspg2*, *Orai*, *OXA1L*, *Dnajc9*, *Usp8*, *ARFGEF3*, and *TBC1D13*. Candidate genes were associated with functions related to neuropeptide signalling, calcium transport, cell adhesion and cytoskeletal organization, mitochondrial function, protein quality control, and intracellular trafficking or signalling. Two candidate regions overlapped between the *XtX* and *C_2_* scans, both on scaffold ptg000103 and associated with *Masc* and *TBC1D13*, respectively (Fig. 2C). The overlapping intervals spanned 33.61 kb and 5.40 kb. Six POD99.9-supported SNPs were shared between the two scans, more than expected by chance (*P* = 2.02 x 10^-14^), indicating strong exact-SNP convergence.

Because host categories were spatially structured across the sampling range, we evaluated the robustness of *C_2_*results using a geographically restricted analysis comprising 16 populations from provinces in which both host categories were represented, including eight populations sampled on *O. ficus-indica* and eight on native *Opuntia* (Fig. 1A). After restricting the dataset to loci that remained polymorphic across these populations, 130,732 SNPs were retained for the sensitivity analysis. The restricted analysis identified 16 candidate regions, compared with 17 in the full-range analysis (Fig. S7). Genome-wide SNP-level *C_2_*statistics showed moderate concordance between analyses (Spearman’s ρ = 0.434; Pearson’s *r* = 0.576), indicating that restricting the geographic composition of the dataset altered the relative strength of many genome-wide signals (Fig. S7). Nevertheless, several of the strongest candidates persisted; 12 of the 17 peak SNPs from the full-range analysis remained within the upper 1% of *C_2_* values in the restricted analysis, including three within the upper 0.1%. Five of the 17 full-range candidate regions physically overlapped candidate regions recovered in the restricted analysis, and four peak-associated genes were shared between analyses. Notably, the *Hspg2* candidate region remained among the strongest signals in the geographically restricted *C_2_* analysis.

### Genotype-environment associations

GEA analyses identified 207 SNP-covariate associations exceeding BF > 20 dB, which clustered into 167 covariate-specific candidate loci across the six environmental and spatial covariates (Fig. S8; Tables S8-S9). Comparison with the differentiation scans revealed limited but notable convergence (Fig. 3A). Three TemperaturePC1-associated GEA loci overlapped a single *XtX* candidate region on scaffold ptg000084, including two loci associated with *fax* and one with *Bag6*. At the SNP level, one *fax*-associated SNP and the *Bag6*-associated SNP were also POD99.9-supported *XtX* SNPs, indicating exact SNP-level convergence. This overlap was significantly greater than expected by chance (*P* = 1.03 x 10^-4^). *Bag6* was the only gene directly shared between the *XtX* and GEA candidate-gene sets (Fig. 2C), although additional TemperaturePC1 loci, including *fax*, showed physical and exact-SNP overlap with an *XtX* candidate region without being the peak-associated gene assigned to that region. No GEA locus physically overlapped a *C_2_*region, although *Hspg2* was independently recovered by the *C_2_* host-use analysis and by the aridity-index and longitude GEAs, representing gene-level convergence without interval-level overlap. The *Hspg2 C_2_* signal also remained among the strongest candidates in the geographically restricted host-use analysis. No candidate gene was shared among *XtX*, *C_2_*, and GEA.

**Figure 3.**
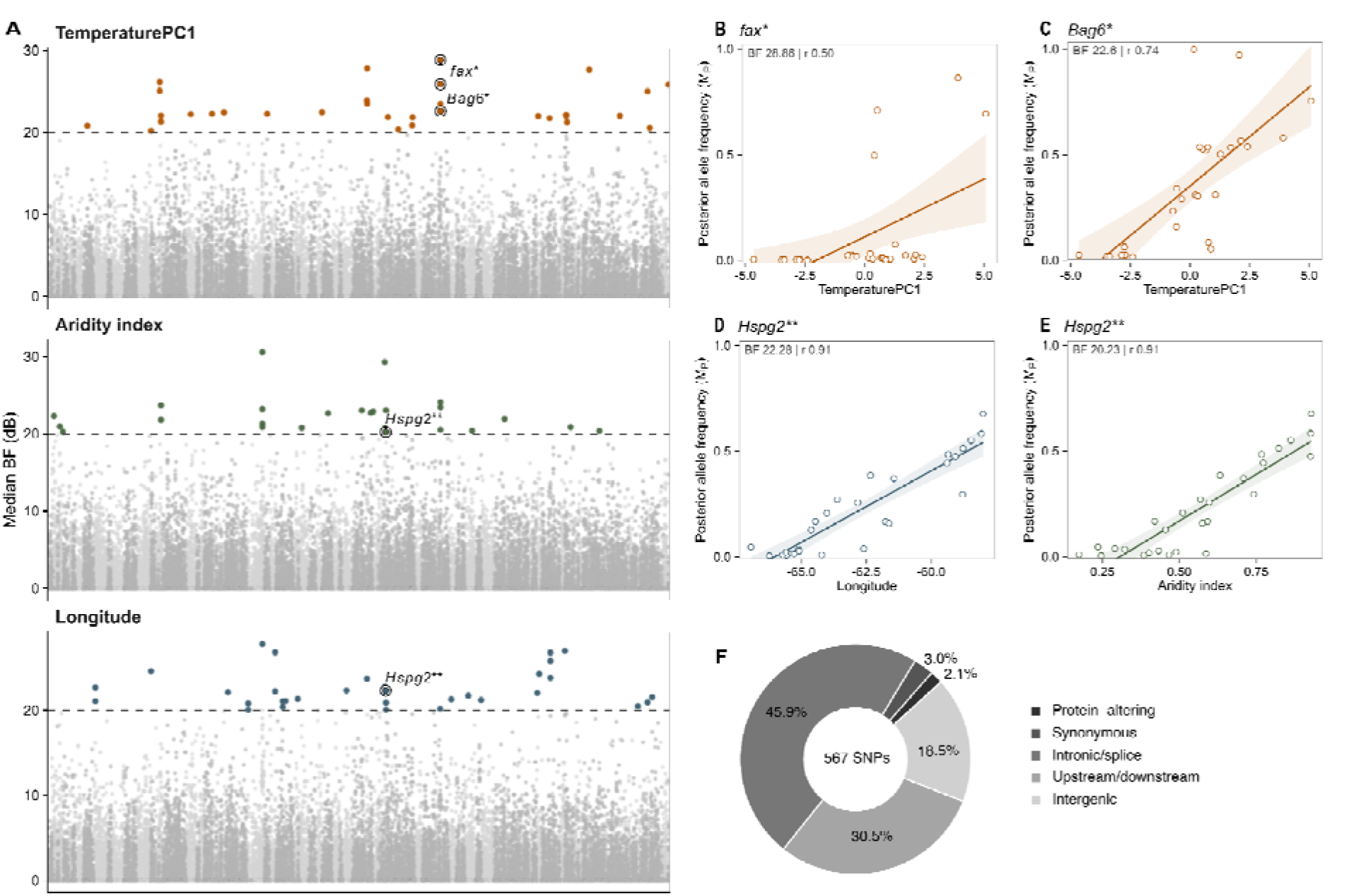
Genotype-environment associations at loci showing convergence with differentiation scans and predicted functional consequences of candidate SNPs. **(A)** Focused BayPass genotype– environment association scans for TemperaturePC1, aridity index, and longitude. SNP-wise association strength is shown as the median Bayes Factors (BFs) across three independent runs, expressed in decibans [BF(dB)]. Colored points indicate SNPs with median BF > 20 dB, and the dashed horizontal line marks this threshold. Highlighted loci correspond to candidates showing convergence with *XtX* or *C_2_* analyses. A single asterisk (*) denotes a GEA locus physically overlapping an *XtX* candidate region, whereas two asterisks (**) denote a GEA candidate associated with a gene also identified by the host-use *C_2_* scan; the latter reflects gene-level convergence and does not imply physical overlap of the underlying candidate intervals. (B-E) BayPass posterior population allele frequencies (M_P_) plotted against the corresponding environmental covariate for **(B)** *fax*-TemperaturePC1, **(C)** *Bag6*-TemperaturePC1, **(D)** *Hspg2*-longitude, and **(E)** *Hspg2*-aridity index. Lines show linear trends with 95% confidence intervals, and inset values report the median BF and descriptive Pearson correlation coefficient (*r*). The same *Hspg2*-associated SNP was recovered for both aridity index and longitude; because these covariates were strongly correlated, the two associations are interpreted as reflecting a shared environmental-geographic signal rather than independent responses to each variable. **(F)** Distribution of predicted functional consequences among the 567 unique candidate SNP positions identified across *XtX*, *C_2_*, and GEA analyses. Each SNP was assigned a single primary consequence according to a hierarchical classification prioritizing protein-altering, UTR, synonymous, non-coding exon/transcript, intronic or splice-region, upstream/downstream, and intergenic consequences. UTR and non-coding exon/transcript classes contained no candidate SNPs and are therefore not displayed.

Allele-frequency clines across environmental gradients were consistent with these associations (Fig. 3B-E). Posterior allele frequency increased moderately with TemperaturePC1 for the strongest *fax* candidate SNP (*r* = 0.50), and the *Bag6* candidate SNP (*r* = 0.74). The *Hspg2* candidate showed strong correlation with aridity index and longitude (*r* = 0.91 for both). Since aridity index and longitude were strongly correlated, these associations were not interpreted as independent environmental effects.

The number of GEA loci varied among covariates, with elevation yielding the most candidates (52), followed by TemperaturePC1 (29), longitude (28), PrecipitationPC1 (26), aridity index (20), and latitude (12) (Table S9). Candidate genes within individual covariates included loci associated with protein homeostasis and neural development (*Bag6*, *fax*) for TemperaturePC1, detoxification and metabolism (*CYP6B2*) for PrecipitationPC1 and elevation, oxygen-sensing pathways (*Egln1*) for aridity index, and circadian regulation (*TIMELESS*) for elevation. Most loci were covariate-specific, although some recurrence occurred among environmental gradients, 26 SNPs exceeded the BF threshold for more than one covariate and 16 candidate genes were recovered across multiple covariates (Fig. S9; Table S10). Sharing was strongest between PrecipitationPC1 and aridity index, and between PrecipitationPC1 and longitude, whereas elevation and latitude showed comparatively little overlap with other covariates. Recurrent genes included *CYP6B2*, shared among PrecipitationPC1, elevation, and longitude, *Hspg2* between aridity index and longitude, and *Pxn* between TemperaturePC1 and aridity index. Representative GEA-only loci unique to each covariate were selected to illustrate environmental associations of candidate SNPs (Fig. S10; Table S11).

### Functional consequences and functional enrichment

Functional annotation of the 567 unique candidate SNP positions identified with *XtX*, *C_2_*, and GEA showed that most candidate variation occurred in non-coding genomic contexts (Fig. 3F). Intronic or splice-region variants were the most frequent class (260 SNPs; 45.9%), followed by upstream/downstream (173; 30.5%) and intergenic variants (105; 18.5%). Only 12 SNPs (2.1%) were predicted to alter protein sequence, while 17 (3.0%) were synonymous variants.

The distribution was broadly similar for the *XtX* and *C_2_* candidate sets, intronic or splice-region variants accounted for 54.1% and 47.1% of candidate SNPs, respectively, whereas GEA candidates were more evenly distributed among intronic/splice-region (32.6%), upstream/downstream (29.1%), and intergenic (33.1%) classes. Overall, protein-altering variants were uncommon, consistent with a substantial contribution of non-coding variation to the genomic signatures detected here. Functional enrichment analyses detected no significantly overrepresented GO terms after FDR correction in the *XtX*, *C_2_*, pooled GEA, covariate-specific GEA, or combined candidate-gene sets (all FDR > 0.05; Table S12).

## Discussion

Our results reveal a heterogeneous genomic landscape of ecological differentiation across the native range of *C. cactorum*, in which different genomic regions were associated with different ecological factors, including climate, geography, and population-level use of the anthropogenically introduced host *O. ficus-indica*. Most candidate regions were specific to individual scans or environmental gradients, suggesting that no single ecological axis or genomic response dominates across the sampled range. Instead, the strongest evidence for local adaptation arose from a limited set of regions supported by complementary analyses. In particular, several temperature-associated loci occurred within regions of elevated population differentiation, while *Hspg2* was recovered in analyses testing for both host use and other environmental factors and remained among the strongest *C_2_* signals after geographically restricting the contrast between host categories. Although convergence among scans was overall limited, exact SNP sharing between some candidate sets was greater than expected by chance, providing additional support for a subset of recurrent genomic signals. Together, these patterns support heterogeneity among genomic regions in their associations with different ecological factors, including both environmental gradients and host use. Moreover, our results suggest that relatively recent anthropogenic changes in host availability (the introduction of *O. ficus-indica*) have become part of the ecological landscape in which native populations of *C. cactorum* differentiate.

The contemporary signals detected here occur against a genomic background shaped by much older geographic and demographic processes. Previous population genomic studies revealed strong differentiation among major geographic lineages and the legacy of Quaternary changes in climate and host distributions, however, contemporary host use explained comparatively little of such population structure (Poveda-Martínez et al., 2023; Andraca-Gómez et al., 2024). Our present results, therefore, not only refine the historical picture consisting of a broad population structure mainly dominated by geography and demographic history, but also shows the effect of present-day ecological associations on a comparatively restricted fraction of the genome. This distinction is important because it suggests that contemporary ecological differentiation can emerge within a strongly structured species without generating a reorganization of genome-wide ancestry. Thus, historical population structure and recent ecological differentiation should therefore be viewed as overlapping temporal layers of genomic variation.

Host-associated differentiation provides an example of the more recent and localized ecological factors affecting genomic variation. In effect, *O. ficus-indica* became part of the South American host landscape only within the last several centuries and is now a widespread crop within the native range of *C. cactorum* (Varone et al., 2014). The moth, nevertheless, remains oligophagous since it uses several *Opuntia* species. However, previous population-genetic studies failed to identify contemporary host use as a major determinant of broad genomic structure. We, therefore, do not interpret the *C_2_* contrast as evidence for genome-wide divergence between host-associated lineages or for an *O. ficus-indica*-specialized host race. Instead, we propose that candidates identified herein point to localized genomic responses without strong neutral divergence or reproductive isolation between moths using different hosts.

This is consistent with the complex nature of host use in herbivorous insects, which involves host recognition, feeding, nutrient acquisition, responses to plant defenses, development, and reproductive behavior. In fact, shifts in host use may therefore involve multiple traits and genomic regions (Simon et al., 2015; Vertacnik and Linnen, 2017). Notably, several of the strongest *C_2_* signals persisted when the analysis was restricted to geographic regions in which both host categories are represented. Although genome-wide concordance between the full and restricted scans was only moderate, most primary peak SNPs remained highly ranked and several candidate regions and genes were recovered independently as candidates. Such persistence suggests that part of the *C_2_* signal reflects localized host-associated differentiation rather than simply reproducing broad geographic divergence, while the incomplete concordance also shows that geographic composition contributes to some of the detected signals. Notably, two host-associated candidate regions were also recovered in the *XtX* differentiation scan. Moreover, exact SNP overlap between *XtX* and *C_2_* candidate sets was greater than expected by chance. This convergence provides additional support for a subset of host-associated regions showing unusually strong differentiation across the native range.

This host-associated signal is especially interesting in light of previous demographic and life-history evidence. Demographic analyses suggested that the population expansion of *C. cactorum* was temporally associated with the establishment and spread of *O. ficus-indica* in Argentina (Poveda-Martínez et al., 2024). The introduced host may therefore have represented not only an increase in host availability and connectivity, but also a resource capable of supporting comparatively high moth performance and, potentially, population growth. Indeed, experimental studies showed that *C. cactorum* generally performed better on cultivated *O. ficus-indica* than on several native South American hosts. Moths reared on *O. ficus-indica* exhibited higher larval survival, shorter development time, and greater potential fecundity (Varone et al., 2012). Such differences could have important demographic consequences, since host-plant quality influences development, survival, and reproduction in herbivorous insects and can thereby affect population-level performance (Awmack & Leather, 2002; Clissold et al., 2015). In Argentina, *C. cactorum* completes three overlapping generations during an approximately nine-month period of activity, followed by winter quiescence at the final larval instar or pupal stage (Folgarait et al., 2018). Together, this evidence suggests that the introduction and establishment of *O. ficus-indica* may have influenced native populations by increasing resource availability and providing favorable developmental and demographic conditions. Our data cannot determine whether the candidate alleles arose after the arrival of *O. ficus-indica* or were already segregating as standing variation that provided the raw material on which selection associated with this relatively recent ecological change could act.

Among the host-associated candidates, *Hspg2* deserves particular attention because it was recovered in several complementary analyses. An intronic region in *Hspg2* was identified in the primary *C_2_* scan, the gene was also associated with aridity and longitude in the GEA analyses, and its *C_2_* signal remained among the strongest candidates after geographic restriction. In vertebrates, *HSPG2* encodes Perlecan, a large heparan-sulfate proteoglycan of the extracellular matrix and basement membranes. In *Drosophila melanogaster*, the orthologous gene is terribly reduced optic lobes (*trol*), which encodes the fly Perlecan homolog (Park et al., 2003; Lindner et al., 2007; Grigorian et al., 2013). Experimental studies in *Drosophila* have shown that *trol* modulates fibroblast growth factor (FGF) and Hedgehog signalling and regulates neuroblast proliferation during development (Park et al., 2003; Lindner et al., 2007). These functions make *Hspg2* an interesting candidate for further study, as developmental signalling and extracellular-matrix organization may influence larval growth and physiological responses to alternative hosts and/or environmental conditions. The host-associated and environmental signals occur in different portions of *Hspg2*, suggesting that distinct variants within the same gene may contribute to associations with host use and environmental variation. Because aridity and longitude are strongly correlated across the sampling range, however, their individual contributions to the environmental association cannot be disentangled. Even so, the repeated recovery of *Hspg2* in independent analyses, along with the persistence of the *C_2_* signal after geographic restriction, makes this locus one of the most compelling candidates identified in the study. We therefore consider *Hspg2* a high-priority target for functional and experimental validation, while avoiding a direct causal interpretation linking it specifically to host use or aridity.

Complementary support for local adaptation came from temperature-associated variation. Three loci associated with TemperaturePC1 overlapped two candidate regions detected by *XtX* analysis, including regions associated with *fax* and *Bag6*. Such convergence is remarkable since GEA and differentiation scans capture different genomic signatures. GEA identifies allele-frequency changes that track an environmental gradient, whereas differentiation scans identify loci that depart from the genome-wide pattern of population divergence (Lotterhos and Whitlock, 2015; Forester et al., 2018). Moreover, exact SNP overlap between candidate regions detected in the TemperaturePC1 GEA and in *XtX* analysis was significantly greater than expected by chance, strengthening the evidence that such convergence is not simply a consequence of incidental genomic overlap. The biological functions of the underlying genes further make these regions interesting candidates. In *Drosophila, fax* (failed axon connections) is expressed in embryonic mesoderm and axons of the central nervous system and interacts genetically with the Abl tyrosine-kinase pathway during axonal development (Hill et al., 1995). *Bag6*, in turn, is involved in protein quality control, membrane-protein biogenesis, ubiquitin-mediated degradation, and regulation of apoptosis, consistent with roles in proteostasis, protein turnover, and cell survival (Lee and Ye, 2013). Although these annotations do not by themselves demonstrate a thermal mechanism, the repeated identification of these regions by both TemperaturePC1 association and *XtX* differentiation provides some of the strongest candidate evidence in our dataset for spatially varying selection. In the absence of direct fitness measurements, *fax* and *Bag6* remain candidates for local adaptation and priority targets for future functional validation.

The wider GEA landscape reinforces a multidimensional view of climatic adaptation. Elevation yielded the largest number of candidate loci, and precipitation, aridity, and longitude shared several signals consistent with their spatial covariation, whereas TemperaturePC1 and latitude contributed comparatively distinct candidate sets. Genes such as *CYP6B2*, a cytochrome P450 implicated in xenobiotic detoxification (Chen et al., 2019), *Egln1*, a core component of the oxygen-sensing pathway (Ivan and Kaelin, 2017), and *TIMELESS*, a circadian gene previously linked to diapause-related adaptation in butterflies (Pruisscher et al., 2018), add plausible functional context to specific covariates, although the correlated nature of the underlying environmental variables precludes the identification of a single potential selective agent. Comparable multidimensional climatic signatures have been reported across independent elevational transects in *Heliconius* (Montejo-Kovacevich et al., 2022), in genotype– environment analyses of the clouded sulfur butterfly (Durkee et al., 2026), and in climate-associated differentiation in the diamondback moth (Chen et al., 2021), suggesting that distributed, multi-axis genomic responses to climate may be a general feature of lepidopterans in heterogeneous landscapes.

Most candidate variants occurred in intronic, intergenic, or gene-proximal regions, whereas predicted protein-altering variants were uncommon. Such non-coding variation may influence gene regulation or be linked to causal variants not directly captured by ddRADseq, consistent with the broader importance of regulatory changes in phenotypic evolution (Wray, 2007; Stern and Orgogozo, 2008). No GO term was significantly enriched after FDR correction, although the relatively small size of some candidate-gene sets may have limited statistical power. Together with the functional diversity of individual candidates, these results are consistent with a heterogeneous genomic response and a potentially polygenic basis of adaptation. Reduced genomic coverage and the absence of direct phenotype or fitness measurements further limit causal inference. Future work combining whole-genome resequencing with functional and ecological validation will be needed to link candidate variants to gene regulation, phenotype, and fitness under contrasting host and environmental conditions.

More broadly, our findings add a contemporary ecological layer to the evolutionary history previously reconstructed for *C. cactorum*. Quaternary climatic change and shifts in native host distributions contributed strongly to the geographic structure that persists across the native range, whereas the recent introduction and spread of *O. ficus-indica* created a widespread new resource in the pre-existing genomic landscape. Our results suggest that contemporary climatic heterogeneity and host use are now associated with differentiation in a more restricted subset of genomic regions, adding recent ecological responses to a deeper demographic background. These results also carry a specific implication for invasion biology. Under the anthropogenically induced adaptation to invade hypothesis (Hufbauer et al., 2012; Estoup et al., 2026), ecological change within the native range of a species, including the incorporation of a widespread new host, could precondition source populations for establishment elsewhere. Whether the host- and climate-associated variants identified here were among those carried into Australia, South Africa, the Caribbean, or North America during subsequent introductions remains untested. Direct comparisons between native and invasive populations at these loci, particularly *Hspg2, fax,* and Bag6, offer a tractable next step and could help determine whether native-range ecological differentiation contributed to the evolutionary potential of *C. cactorum* as an invasive species.

## Author contribution

DPM: Conceptualization, Methodology, Formal analysis, Visualization, Supervision, Writing - original draft. MNJP: Methodology, Data curation, Formal analysis, Writing - review & editing. NAS: Methodology, Writing - review & editing. NNM: Writing - review & editing. EH: Supervision - Writing - review & editing. LV: Supervision, Writing - review & editing. All authors read and approved the final version of the manuscript.

## Acknowledgements

We would like to thank Mariel Guala and Malena Fuentes Corona for fieldwork support. Computational resources were provided by the Genotoul-Bioinfo facility, which we gratefully acknowledge.

## Ethics declarations

The manuscript presents research on animals that do not require ethical approval for their study.

## Data availability statement

All data used in this study are described in the main article and Supplementary Material. Sampling localities and associated metadata are provided in Table S1. The ddRAD-seq data are available from the NCBI Sequence Read Archive under BioProject PRJNA666743. The all-SNP and unlinked-SNP datasets generated for this study as well as BayPass input files were archived in the Figshare Digital Repository (Poveda-Martínez and Parany, 2026) https://doi.org/10.6084/m9.figshare.33340938.

## Supporting information

Supplementary Information

Supplementary tables with metadata spreadsheet-based tables with 12 tabs, each containing additional information to Supplementary Table S1 - S12.

---

**Table 1.**
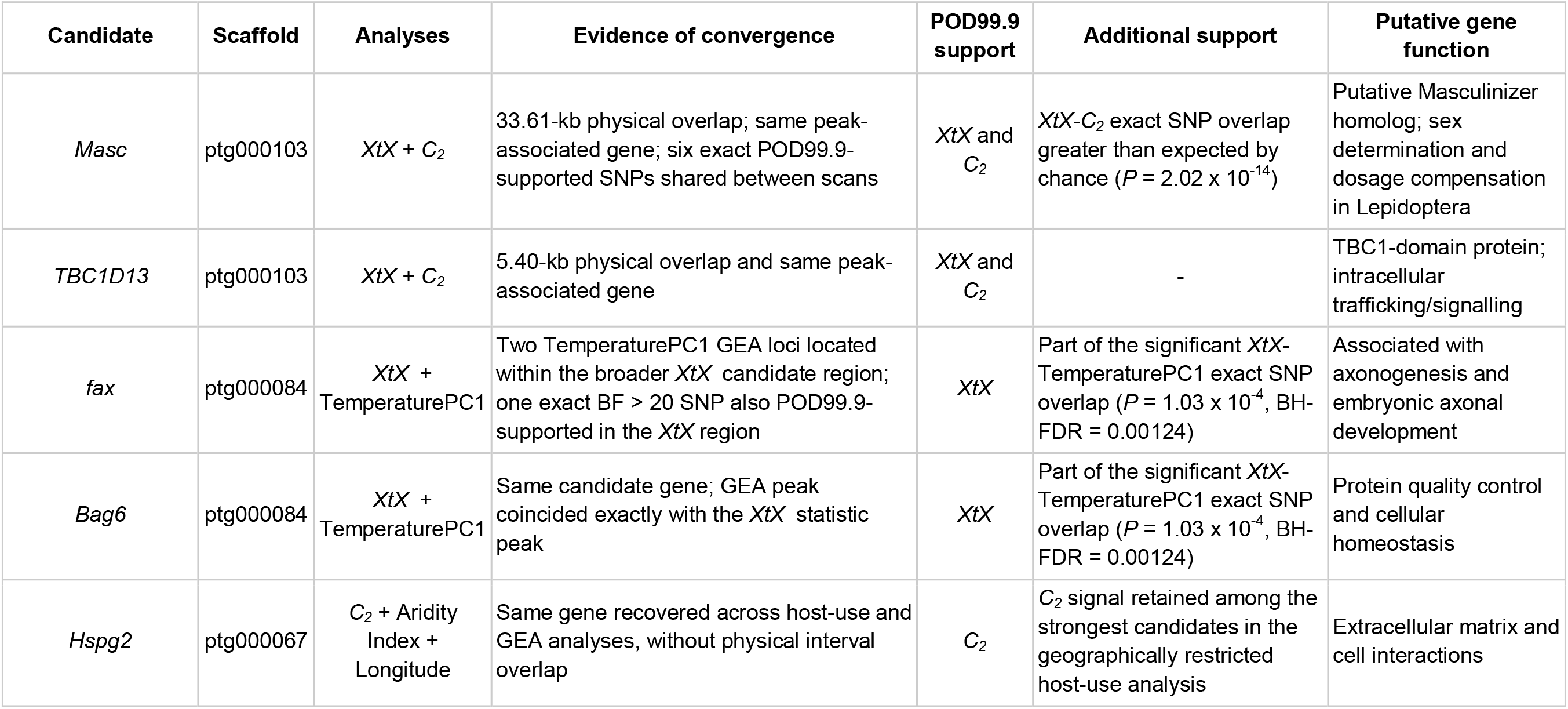
Prioritized candidate loci associated with putative local adaptation in *Cactoblastis cactorum*. *XtX*, BayPass genome-wide differentiation statistic; *C_2_*, host-use contrast; GEA, genotype-environment association; BF, Bayes factor. Candidates were prioritized based on recurrence across analyses at the SNP, genomic-region, or gene level, and/or additional support from the geographically restricted host-use analysis. POD99.9 indicates SNPs exceeding the 99.9th percentile of the model-based neutral posterior predictive distribution.

## References

Alexa A, Rahnenführer J (2026). topGO: Enrichment Analysis for Gene Ontology. doi:10.18129/B9.bioc.topGO. R package version 2.64.0, https://bioconductor.org/packages/topGO.

Alexander DH, Novembre J, Lange K. (2009). Fast model-based estimation of ancestry in unrelated individuals. Genome Res;19(9):1655–64.

Andraca-Gómez, G., Ordano, M., Lira-Noriega, A., Osorio-Olvera, L., Domínguez, C. A., & Fornoni, J. (2024). Climatic and soil characteristics account for the genetic structure of the invasive cactus moth *Cactoblastis cactorum*, in its native range in Argentina. PeerJ, 12, e16861.

Awmack, C. S., & Leather, S. R. (2002). Host plant quality and fecundity in herbivorous insects. Annual Review of Entomology, 47, 817–844

Bloem S, Mizell III RF, Bloem KA, Hight SD, Carpenter JE. Laboratory evaluation of insecticides for control of the invasive *Cactoblastis cactorum* (Lepidoptera: Pyralidae). Florida Entomologist. 2005;88:395–400.

Brooks, C. P., Ervin, G. N., Varone, L., & Logarzo, G. A. (2012). Native ecotypic variation and the role of host identity in the spread of an invasive herbivore, *Cactoblastis cactorum*. Ecology, 93(2), 402–410.

Camacho, C., Coulouris, G., Avagyan, V., Ma, N., Papadopoulos, J., Bealer, K., & Madden, T. L. (2009). BLAST+: architecture and applications. BMC Bioinformatics, 10, 421.

Carroll, S. P., & Boyd, C. (1992). Host race radiation in the soapberry bug: natural history with the history. Evolution, 46(4), 1052–1069.

Carroll, S. P., Dingle, H., & Klassen, S. P. (1997). Genetic differentiation of fitness-associated traits among rapidly evolving populations of the soapberry bug. Evolution, 51(4), 1182–1188.

Chang, C. C., Chow, C. C., Tellier, L. C. A. M., Vattikuti, S., Purcell, S. M., & Lee, J. J. (2015). Second-generation PLINK: rising to the challenge of larger and richer datasets. GigaScience, 4, 7.

Chen, Y., Liu, Z., Régnière, J., Vasseur, L., Lin, J., Huang, S., Ke, F., Chen, S., Li, J., Huang, J., Gurr, G. M., You, M., & You, S. (2021). Large-scale genome-wide study reveals climate adaptive variability in a cosmopolitan pest. Nature Communications, 12, 7206.

Chen, S., Elzaki, M. E. A., Ding, C., Li, Z.-F., Wang, J., Zeng, R.-S., & Song, Y.-Y. (2019). Plant allelochemicals affect tolerance of polyphagous lepidopteran pest *Helicoverpa armigera* (Hübner) against insecticides. Pesticide Biochemistry and Physiology, 154, 32–38.

Clissold, F. J., Coggan, N., & Simpson, S. J. (2015). Temperature, food quality and life history traits of herbivorous insects. Current Opinion in Insect Science, 11, 63–70.

Danecek, P., Auton, A., Abecasis, G., Albers, C. A., Banks, E., DePristo, M. A., Handsaker, R. E., Lunter, G., Marth, G. T., Sherry, S. T., McVean, G., & Durbin, R. (2011). The variant call format and VCFtools. Bioinformatics, 27(15), 2156–2158.

Danecek, P., Bonfield, J. K., Liddle, J., Marshall, J., Ohan, V., Pollard, M. O., Whitwham, A., Keane, T., McCarthy, S. A., Davies, R. M., & Li, H. (2021). Twelve years of SAMtools and BCFtools. GigaScience, 10(2), giab008

Dlugosch, K. M., & Parker, I. M. (2008). Founding events in species invasions: Genetic variation, adaptive evolution, and the role of multiple introductions. Molecular Ecology, 17(1), 431–449.

Dlugosch, K. M., Anderson, S. R., Braasch, J., Cang, F. A., & Gillette, H. D. (2015). The devil is in the details: Genetic variation in introduced populations and its contributions to invasion. Molecular Ecology, 24(9), 2095–2111.

Durkee, L. F., Bossu, C. M., Ruegg, K. C., Forester, B. R., Opler, P. A., & Hufbauer, R. A. (2026). Summer rainfall drives adaptation with gene flow in a widespread butterfly. Molecular Ecology, 35(12), e70388.

Estoup, A., Hulme, P. E., Hodgins, K. A., Gautier, M., Lombaert, E., Dhami, M. K., McGaughran, A., & Hufbauer, R. A. (2026). Can habitat modification in the native range promote invasion? Trends in Ecology & Evolution. Advance online publication. 10.1016/j.tree.2026.06.006

Estoup, A., Ravigné, V., Hufbauer, R. A., Vitalis, R., Gautier, M., & Facon, B. (2016). Is there a genetic paradox of biological invasion? Annual Review of Ecology, Evolution, and Systematics, 47, 51–72.

Ewels, P., Magnusson, M., Lundin, S., & Käller, M. (2016). MultiQC: summarize analysis results for multiple tools and samples in a single report. Bioinformatics, 32(19), 3047–3048.

Fariello, M. I., Boitard, S., Mercier, S., Robelin, D., Faraut, T., Arnould, C., & SanCristobal, M. (2017). Accounting for linkage disequilibrium in genome scans for selection without individual genotypes: the local score approach. Molecular ecology, 26(14), 3700–3714.

Fick, S. E., & Hijmans, R. J. (2017). WorldClim 2: new 1 km spatial resolution climate surfaces for global land areas. International journal of climatology, 37(12), 4302–4315.

Folgarait, P. J., Montenegro, G. A., Plowes, R. M., & Gilbert, L. (2018). A study of *Cactoblastis cactorum* (Lepidoptera: Pyralidae) in its native range: further insights into life cycle, larval identification, developmental parameters, natural enemies, and damage to the host plant *Opuntia ficus-indica*. Florida Entomologist, 101(4), 559–572.

Forester, B. R., Lasky, J. R., Wagner, H. H., & Urban, D. L. (2018). Comparing methods for detecting multilocus adaptation with multivariate genotype–environment associations. Molecular Ecology, 27(9), 2215–2233.

Förstner, W., & Moonen, B. (2003). A metric for covariance matrices. In Geodesy-the Challenge of the 3rd Millennium (pp. 299–309). Berlin, Heidelberg: Springer Berlin Heidelberg.

Fuentes Corona, M., Mengoni Goñalons, C., Cecere, M. C., Hight, S. D., Poveda-Martínez, D., & Varone, L. (2025). Impact of cactus moth (Lepidoptera: Pyralidae) pest densities on fruit production and quality in cactus pear. Journal of Economic Entomology, 118(3), 1262–1270.

Fury, W., Batliwalla, F., Gregersen, P. K., & Li, W. (2006). Overlapping probabilities of top ranking gene lists, hypergeometric distribution, and stringency of gene selection criterion. Conference Proceedings of the IEEE Engineering in Medicine and Biology Society, 2006, 5531–5534.

Gautier, M. (2015). Genome-Wide Scan for Adaptive Divergence and Association with Population-Specific Covariates. Genetics, 201(4), 1555–1579.

Griffith, M. P. (2004). The origins of an important cactus crop, *Opuntia ficus-indica* (Cactaceae): New molecular evidence. American Journal of Botany, 91(11), 1915–1921.

Grigorian, M., Liu, T., Banerjee, U., & Hartenstein, V. (2013). The proteoglycan Trol controls the architecture of the extracellular matrix and balances proliferation and differentiation of blood progenitors in the *Drosophila* lymph gland. Developmental Biology, 384(2), 301–312

Hight SD, Carpenter JE, Bloem S, Bloem KA. Developing a sterile insect release program for *Cactoblastis cactorum* (Berg)(Lepidoptera:Pyralidae): Effective overflooding ratios and release-recapture field studies. Environmental entomology. 2005; 34:850–856

Hijmans, R. J., Barbosa, M., Ghosh, A., & Mandel, A. (2026). geodata: Access Geographic Data. R package version 0.6–9. 10.32614/CRAN.package.geodata

Hill, K. K., Bedian, V., Juang, J.-L., & Hoffmann, F. M. (1995). Genetic interactions between the *Drosophila* Abelson (Abl) tyrosine kinase and *failed axon connections* (*fax*), a novel protein in axon bundles. Genetics, 141(2), 595–606.

Hufbauer, R. A., Facon, B., Ravigné, V., Turgeon, J., Foucaud, J., Lee, C. E., Rey, O., & Estoup, A. (2012). Anthropogenically induced adaptation to invade (AIAI): Contemporary adaptation to human-altered habitats within the native range can promote invasions. Evolutionary Applications, 5(1), 89– 101.

Inglese, P., Mondragón, C., Nefzaoui, A., & Sáenz, C. (Eds.). (2017). Crop ecology, cultivation and uses of cactus pear. Rome: Food and Agriculture Organization of the United Nations and International Center for Agricultural Research in the Dry Areas. ISBN 978-92-5-109860-8.

Ivan, M., & Kaelin, W. G. Jr. (2017). The EGLN-HIF O sensing system: Multiple inputs and feedbacks. Molecular Cell, 66(6), 772–779.

Jones, P., Binns, D., Chang, H.-Y., Fraser, M., Li, W., McAnulla, C., et al. (2014). InterProScan 5: genome-scale protein function classification. Bioinformatics, 30(9), 1236–1240.

Kiesling, R. (1988). Cactaceae. In M. N. Correa (Ed.), Flora Patagónica: Dicotiledóneas Dialipétalas (Oxalidaceae a Cornaceae), Vol. 5, pp. 218–243. Instituto Nacional de Tecnología Agropecuaria (INTA), Buenos Aires, Argentina.

Körner, C. (2007). The use of ‘altitude’ in ecological research. Trends in Ecology & Evolution, 22(11), 569–574.

Lee, C. E. (2002). Evolutionary genetics of invasive species. Trends in Ecology & Evolution, 17(8), 386–391.

Lee, J. G., & Ye, Y. (2013). Bag6/Bat3/Scythe: a novel chaperone activity with diverse regulatory functions in protein biogenesis and degradation. Bioessays, 35(4), 377–385.

Lindner, J. R., Hillman, P. R., Barrett, A. L., Jackson, M. C., Perry, T. L., Park, Y., & Datta, S. (2007). The *Drosophila* Perlecan gene trol regulates multiple signaling pathways in different developmental contexts. BMC Developmental Biology, 7, 121.

Lotterhos, K. E., & Whitlock, M. C. (2015). The relative power of genome scans to detect local adaptation depends on sampling design and statistical method. Molecular Ecology, 24(5), 1031– 1046.

Marsico, T. D., Wallace, L. E., Ervin, G. N., Brooks, C. P., McClure, J. E., & Welch, M. E. (2011). Geographic patterns of genetic diversity from the native range of *Cactoblastis cactorum* (Berg) support the documented history of invasion and multiple introductions for invasive populations. Biological Invasions, 13, 857–868.

Matsubayashi, K. W., Ohshima, I., & Nosil, P. (2010). Ecological speciation in phytophagous insects. Entomologia Experimentalis et Applicata, 134(1), 1–27.

McFadyen, R. E. C. (1985). Larval characteristics of *Cactoblastis* spp. (Lepidoptera: Pyralidae) and the selection of species for biological control of prickly pears (Opuntia spp.). Bulletin of Entomological Research, 75(1), 159–168.

McLaren, W., Gil, L., Hunt, S. E., Riat, H. S., Ritchie, G. R. S., Thormann, A., et al. (2016). The Ensembl Variant Effect Predictor. Genome Biology, 17, 122

Montejo-Kovacevich, G., Meier, J. I., Bacquet, C. N., Warren, I. A., Chan, Y. F., Kucka, M., Salazar, C., Rueda-M, N., Montgomery, S. H., McMillan, W. O., Kozak, K. M., Nadeau, N. J., Martin, S. H., & Jiggins, C. D. (2022). Repeated genetic adaptation to altitude in two tropical butterflies. Nature Communications, 13, 4676.

Montenegro, G., Acosta, M.C., Caeiro, L. et al. Tracing the geographic origins of two forms of Opuntia ficus-indica cultivated in Argentina using haplotype diversity patterns, and cytogenetic and morphological analyses. Genet Resour Crop Evol 71, 3915–3930 (2024). 10.1007/s10722-024-01876-w

Morrone J. J. (2014). Biogeographical regionalisation of the Neotropical region. Zootaxa 3782 (1), 1– 110.

Nugroho, A., S. Comte, P. Barrière, R. B. Allen, W. B. Sherwin, and L. A. Rollins. 2026. Two Secondary Introductions From a Shared Bridgehead Population Show Evidence of Divergent and Parallel Selection. Molecular Ecology 35, no. 15: e70495.

Olazcuaga, L., Loiseau, A., Parrinello, H., Paris, M., Fraimout, A., Guedot, C., Diepenbrock, L. M., Kenis, M., Zhang, J., Chen, X., Borowiec, N., Facon, B., Vogt, H., Price, D. K., Vogel, H., Prud’homme, B., Estoup, A., & Gautier, M. (2020). A Whole-Genome Scan for Association with Invasion Success in the Fruit Fly Drosophila suzukii Using Contrasts of Allele Frequencies Corrected for Population Structure. Molecular Biology and Evolution, 37(8), 2369–2385.

Oliveira, E. A. D., Perez, M. F., Bertollo, L. A. C., Gestich, C. C., Ráb, P., Ezaz, T., & Cioffi, M. B. (2020). Historical demography and climate driven distributional changes in a widespread Neotropical freshwater species with high economic importance. Ecography, 43(9), 1291–1304.

Park, Y., Rangel, C., Reynolds, M. M., Caldwell, M. C., Johns, M., Nayak, M., Welsh, C. J. R., McDermott, S., & Datta, S. (2003). *Drosophila* Perlecan modulates FGF and Hedgehog signals to activate neural stem cell division. Developmental Biology, 253(2), 247–257.

Peterson, B. K., Weber, J. N., Kay, E. H., Fisher, H. S., & Hoekstra, H. E. (2012). Double digest RADseq: an inexpensive method for de novo SNP discovery and genotyping in model and non-model species. PloS one, 7(5), e37135.

Poveda-Martínez, D., Moreyra, N.N., Hasson, E. et al. (2024). Demographic inference provides evidence of a quaternary-driven impact on the cactus moth and sheds light on the putative role of an exotic host. Biol Invasions 26, 2313–2327. 10.1007/s10530-024-03317-2

Poveda-Martínez, Daniel; Parany, Mamionah N. J. (2026). Genomic datasets for selection scans and genotype–environment association analyses in *Cactoblastis cactorum*. figshare. Dataset. 10.6084/m9.figshare.33340938.v1

Poveda-Martínez, D., Noguerales, V., Hight, S. D., Logarzo, G., Emerson, B. C., Varone, L., & Hasson, E. (2023). Geography, climate and shifts in host plants distribution explain the genomic variation in the cactus moth. Frontiers in Ecology and Evolution, 11, 1260857.

Pruisscher, P., Nylin, S., Gotthard, K., & Wheat, C. W. (2018). Genetic variation underlying local adaptation of diapause induction along a cline in a butterfly. Molecular Ecology, 27, 3613–3626.

Rochette, N. C., Rivera Colón, A. G., & Catchen, J. M. (2019). Stacks 2: Analytical methods for paired end sequencing improve RADseq based population genomics. Molecular ecology, 28(21), 4737–4754.

Sakai, A. K., Allendorf, F. W., Holt, J. S., Lodge, D. M., Molofsky, J., With, K. A., Baughman, S., Cabin, R. J., Cohen, J. E., Ellstrand, N. C., McCauley, D. E., O’Neil, P., Parker, I. M., Thompson, J. N., & Weller, S. G. (2001). The population biology of invasive species. Annual Review of Ecology and Systematics, 32, 305–332.

Simon, J.-C., d’Alençon, E., Guy, E., Jacquin-Joly, E., Jaquiéry, J., Nouhaud, P., Peccoud, J., Sugio, A., & Streiff, R. (2015). Genomics of adaptation to host-plants in herbivorous insects. Briefings in Functional Genomics, 14(6), 413–423.

Stern, D. L., & Orgogozo, V. (2008). The loci of evolution: How predictable is genetic evolution? Evolution, 62, 2155–2177.

Varone, L., Acosta, M. M., Logarzo, G. A., Briano, J. A., Hight, S. D., & Carpenter, J. E. (2012). Laboratory Performance of *Cactoblastis cactorum* (Lepidoptera: Pyralidae) on South and North American *Opuntia* species occurring in Argentina. Florida Entomologist, 95(4), 1163–1173.

Varone, L., Logarzo, G. A., Briano, J. A., Hight, S. D., & Carpenter, J. E. (2014). *Cactoblastis cactorum* (Berg) (Lepidoptera: Pyralidae) use of *Opuntia* host species in Argentina. Biological Invasions, 16, 2367–2380.

Varone, L., Mengoni Goñalons, C., Faltlhauser, A. C., Guala, M. E., Wolaver, D., Srivastava, M., & Hight, S. D. (2020). Effect of rearing *Cactoblastis cactorum* on an artificial diet on the behaviour of *Apanteles opuntiarum*. Journal of Applied Entomology, 144, 278–286.

Vasimuddin Md, Sanchit Misra, Heng Li, Srinivas Aluru. Efficient Architecture-Aware Acceleration of BWA-MEM for Multicore Systems. IEEE Parallel and Distributed Processing Symposium (IPDPS), 2019. 10.1109/IPDPS.2019.00041

Vertacnik, K. L., & Linnen, C. R. (2017). Evolutionary genetics of host shifts in herbivorous insects: insights from the age of genomics. Annals of the New York Academy of Sciences, 1389(1), 186–212.

Wray, G. A. (2007). The evolutionary significance of cis-regulatory mutations. Nature Reviews Genetics, 8, 206–216.

Zimmermann, H. G., Moran, V. C., & Hoffmann, J. H. (2000). The renowned cactus moth, *Cactoblastis cactorum*: its natural history and threat to native *Opuntia* floras in Mexico and the United States of America. Florida Entomologist, 83(4), 543–551.

Zomer, R. J., Xu, J., & Trabucco, A. (2022). Version 3 of the Global Aridity Index and Potential Evapotranspiration Database. Scientific Data, 9, 409.

