## Supplementary Information for "Contemporary ecological heterogeneity shapes candidate adaptive genomic variation across the native range of an invasive herbivore"

**This PDF file includes:**

Figures S1 to S10.

Additional supplementary material for this work includes:
Supplementary tables with metadata spreadsheet-based tables with 12 tabs, each containing additional information to Supplementary Table S1 - S12.

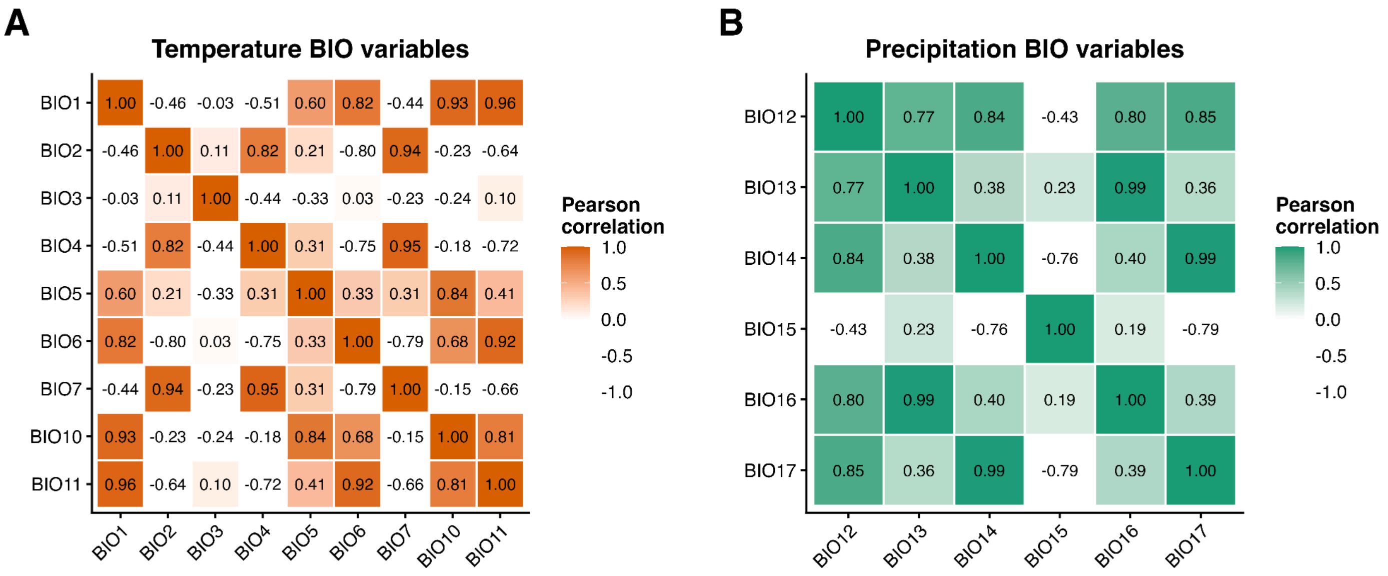


**Figure S1. Correlation structure among climatic variables used for principal component analyses.** Pearson correlation matrices are shown for (A) temperature-related WorldClim bioclimatic variables (BIO1–BIO7, BIO10, and BIO11) and (B) precipitation-related variables (BIO12–BIO17). Cell values indicate Pearson correlation coefficients, with stronger color intensity representing stronger positive correlations. These variables were subsequently summarized separately using principal component analyses to derive the temperature and precipitation climatic axes used in downstream analyses.

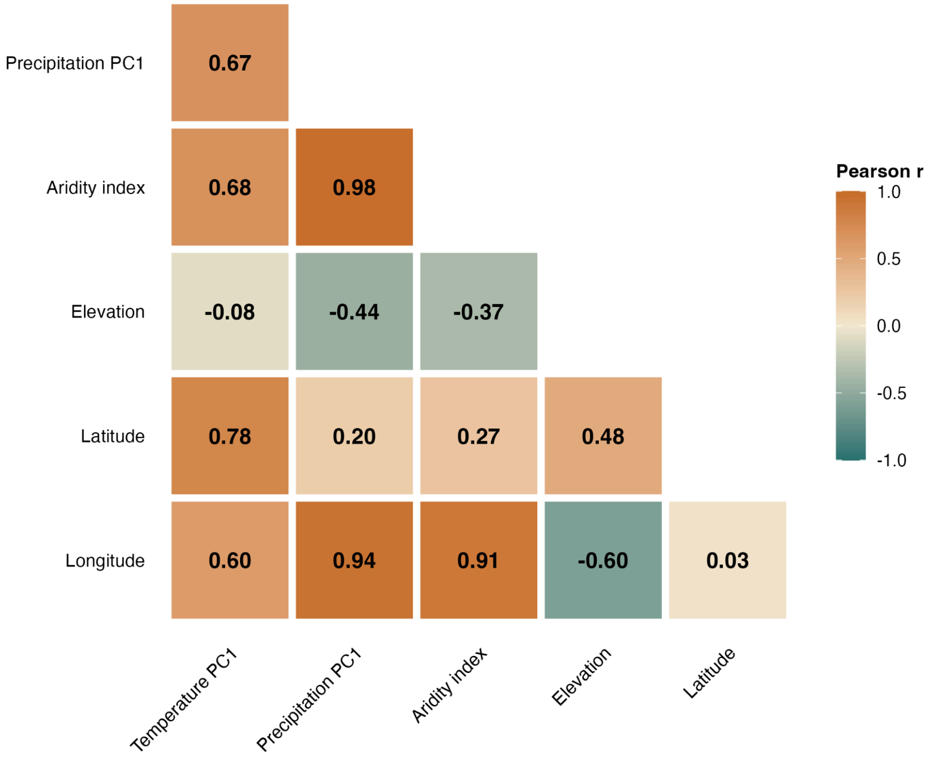


**Figure S2. Pairwise correlations among environmental and spatial covariates used in genotype–environment association analyses.** Pearson correlation coefficients (r) were calculated across the 28 *Cactoblastis cactorum* sampling populations for TemperaturePC1, PrecipitationPC1, aridity index, elevation, latitude, and longitude. Only the lower triangle of the correlation matrix is shown, with cell colors indicating the direction and magnitude of the correlation and numerical values reporting Pearson's r. Several covariates showed strong correlations. Each covariate was nevertheless tested independently in separate BayPass genotype–environment association models.

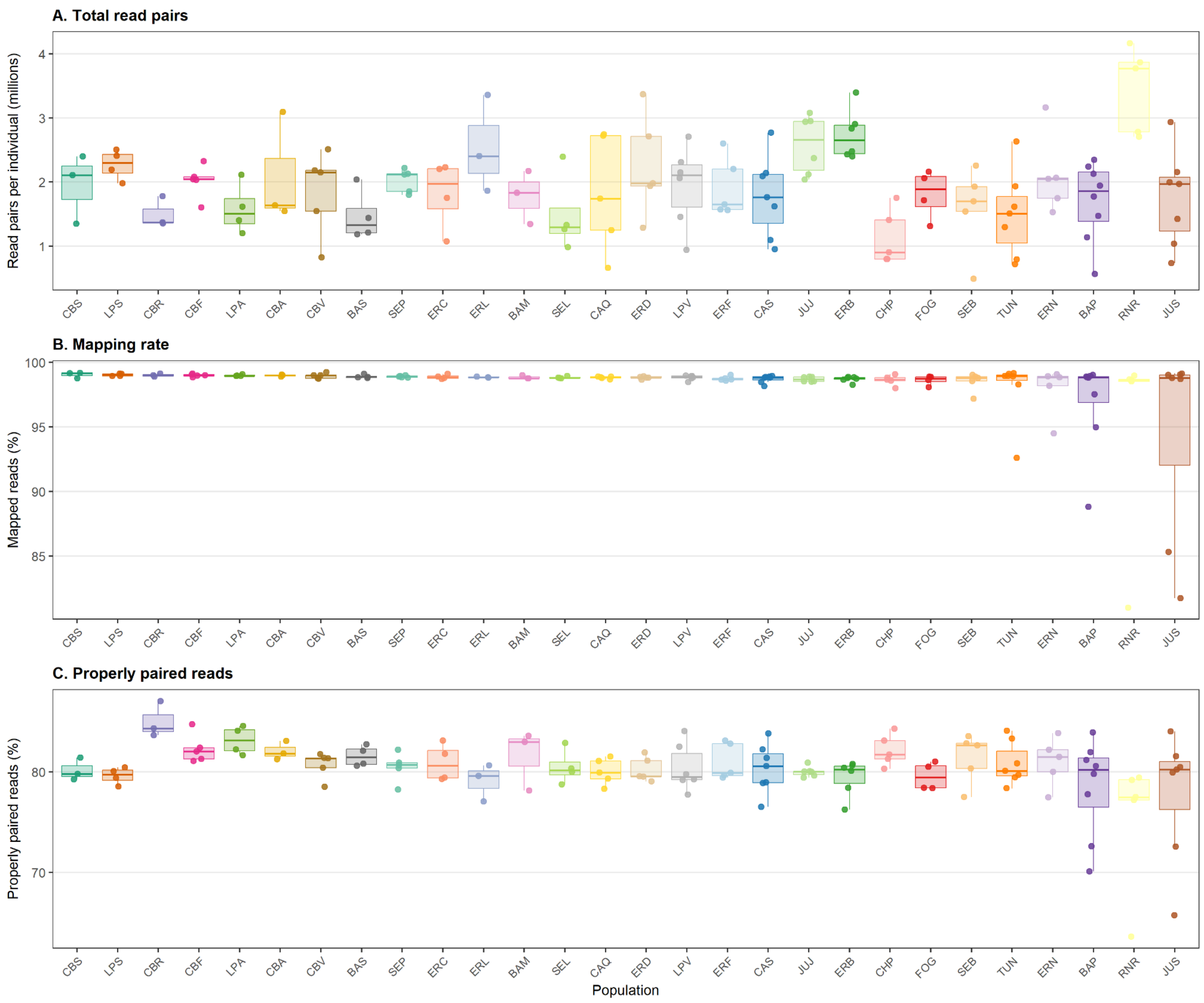


**Figure S3. Mapping statistics across cactus moth ddRADseq samples.** (A) Number of paired-end read pairs per individual after filtering and alignment. (B) Percentage of reads mapped to the *Cactoblastis cactorum* reference genome using bwa-mem2. (C) Percentage of properly paired reads per individual. Points represent individual samples, and boxplots summarize the distribution within each population.

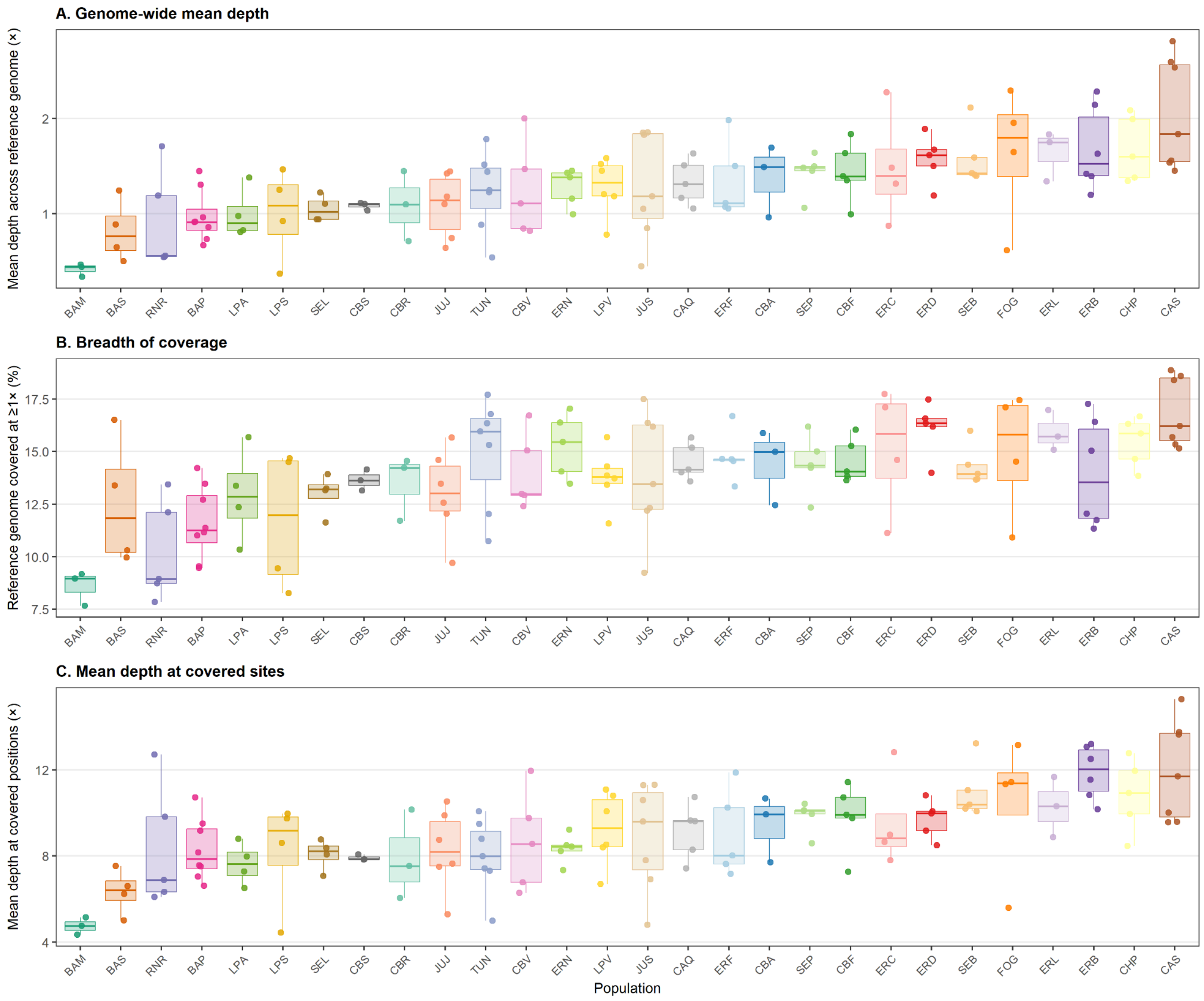


**Figure S4. Coverage statistics across cactus moth ddRADseq samples grouped by population.** (A) Genome-wide mean depth estimated across the full *Cactoblastis cactorum* reference genome. (B) Breadth of coverage, measured as the percentage of the reference genome covered by at least one read. (C) Mean depth at covered genomic positions, calculated only for sites with coverage ≥1×. Points represent individual samples, and boxplots summarize the distribution within each population. Populations are ordered by increasing genome-wide mean depth. The higher values observed for mean depth at covered sites compared with genome-wide mean depth are expected for a reduced-representation ddRADseq dataset, where only a subset of genomic regions is sequenced.

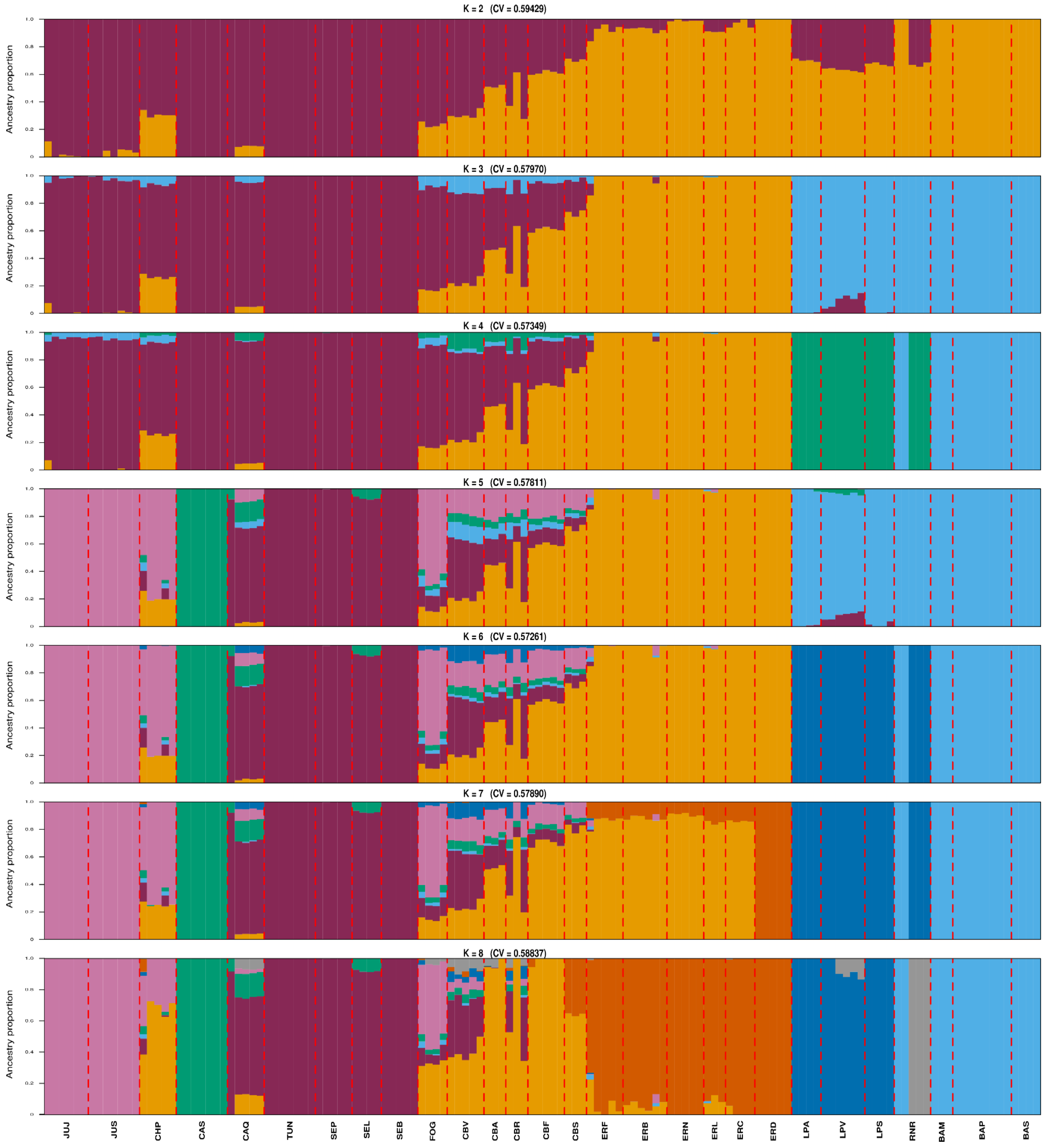


**Figure S5. Ancestry proportions across *K* = 2–8 for *Cactoblastis cactorum* populations sampled across the native range in Argentina.** Each vertical bar represents an individual, and colors indicate inferred ancestry components. The lowest cross-validation error was obtained at (*K* = 6), which revealed finer-scale spatial structure corresponding approximately to northern, central, western, southeastern, southwestern, and eastern groups. At (*K* = 3), the three major geographically structured lineages previously described across the native range were recovered: central–northern–western, eastern, and southern lineages. Admixture was particularly evident among northern and central populations and remained visible across several (*K*) values, whereas no clear clustering according to host plant was observed.

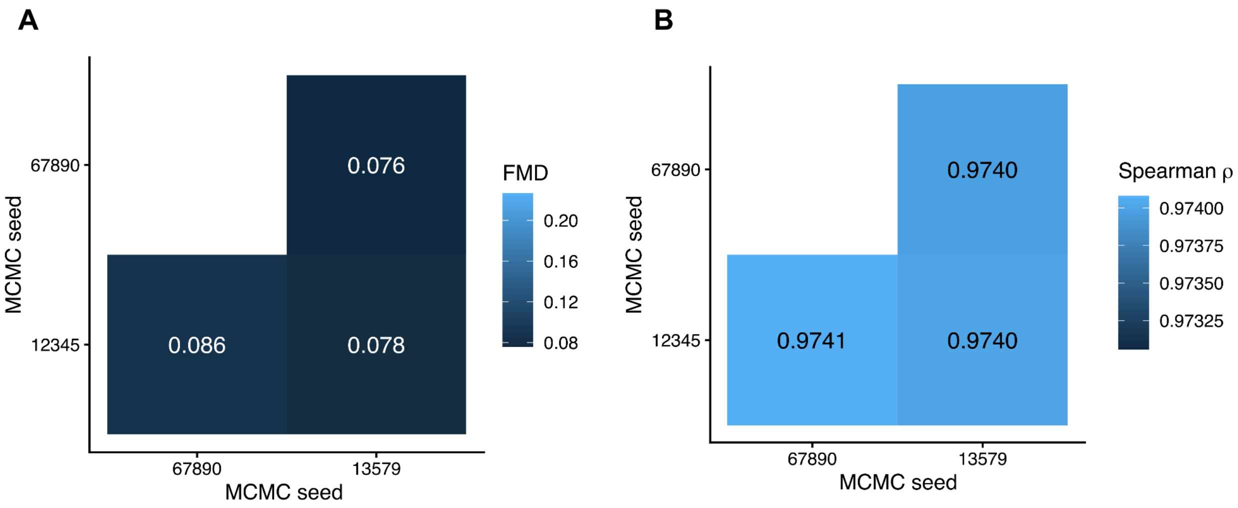


**Figure S6. Convergence and reproducibility of BayPass analyses across independent MCMC chains for the all-SNP dataset.** (A) Pairwise Förstner–Moonen distances (FMD) between population covariance matrices ($\Omega$) inferred from three independent BayPass runs initiated with different random seeds. Lower FMD values indicate greater similarity among $\Omega$ estimates; pairwise FMD values ranged from 0.076 to 0.086. (B) Pairwise Spearman rank correlations of SNP-level *XtX* estimates among the same three independent runs. *XtX* estimates were highly concordant across chains, with Spearman's $\rho$ ranging from 0.9740 to 0.9741. Together, these diagnostics indicate stable estimation of the population covariance matrix and high reproducibility of genome-wide *XtX* estimates across independent MCMC runs for the all-SNP dataset.

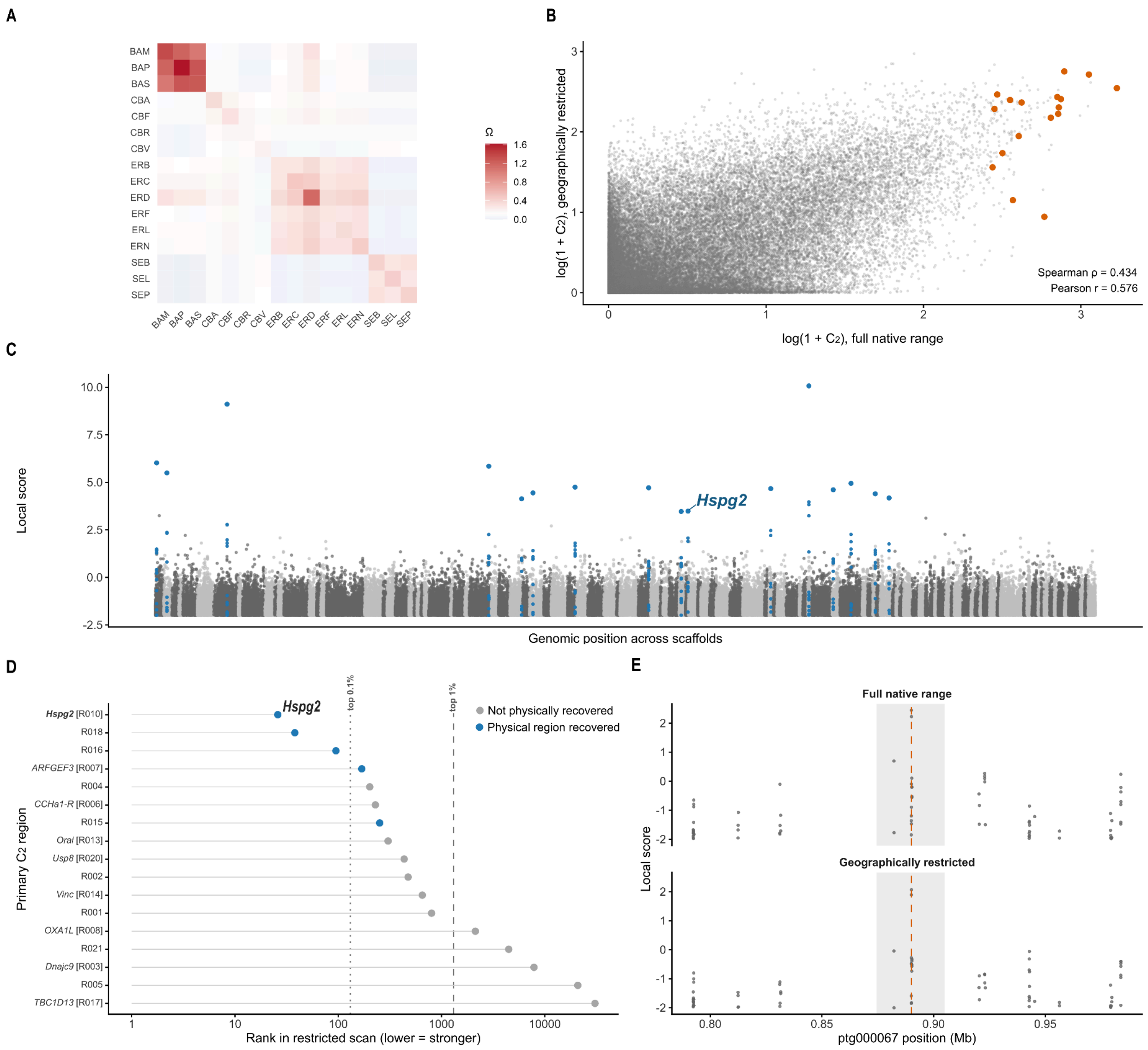


**Figure S7. Geographic sensitivity of host-associated genomic differentiation in *Cactoblastis cactorum*.** The BayPass $C_2$ host-use contrast was repeated using 16 geographically restricted populations representing *O. ficus-indica* and native *Opuntia*. (**A**) Population covariance matrix (Ω) estimated for the restricted dataset. (**B**) Genome-wide correspondence between full-range and restricted $C_2$ statistics for 130,732 shared SNPs; orange points indicate the 17 final primary $C_2$ peak SNPs. (**C**) Restricted local-score genome scan based on scaffolds >1 Mb, with SNPs within the 16 candidate regions highlighted in blue and selected peak-associated genes labelled. (**D**) Restricted-scan ranks of the 17 primary $C_2$ peaks. Blue points indicate primary regions physically recovered in the restricted analysis, whereas grey points indicate regions without physical overlap; 12/17 peaks remained within the top 1% of restricted $C_2$ values, including three within the top 0.1%, and five primary regions were physically recovered. (**E**) Local-score profiles surrounding *Hspg2* in the full-range and restricted analyses. The shaded interval indicates the $C_2$ candidate region, and the orange dashed line marks its full-range statistic peak. Overall, the analysis shows that several of the strongest host-associated signals persisted after geographic restriction, although recovery of only a subset of the full-range candidate regions indicates that some signals were sensitive to the geographic composition of the population contrast.

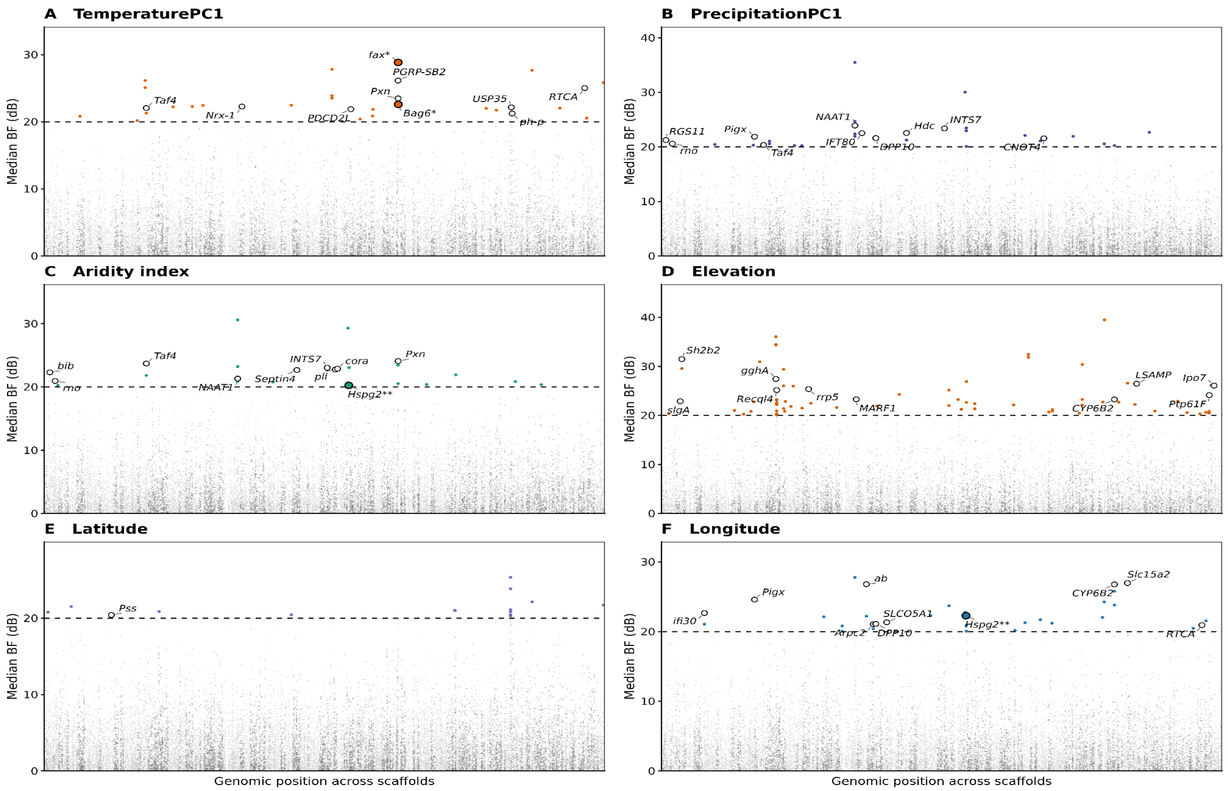


**Figure S8. Genome-wide genotype–environment association scans across six environmental and spatial covariates**. BayPass results are shown for (A) TemperaturePC1, (B) PrecipitationPC1, (C) aridity index, (D) elevation, (E) latitude, and (F) longitude. Association strength corresponds to the median Bayes Factor across three independent runs, expressed in decibans [BF(dB)]. SNPs with median BF > 20 dB are highlighted, and the dashed horizontal line marks this threshold. Candidate SNPs on the same scaffold and separated by ≤1 kb were grouped into loci, with the highest-BF SNP defining the locus peak. Up to 10 biologically annotated candidate genes are labelled per panel, prioritizing cross-analysis candidates and then the strongest associations. A single asterisk (*) indicates a GEA locus physically overlapping an *XtX* candidate region, whereas two asterisks (**) indicate a gene also identified by the host-use $C_2$ analysis, without necessarily implying physical overlap of the underlying intervals.

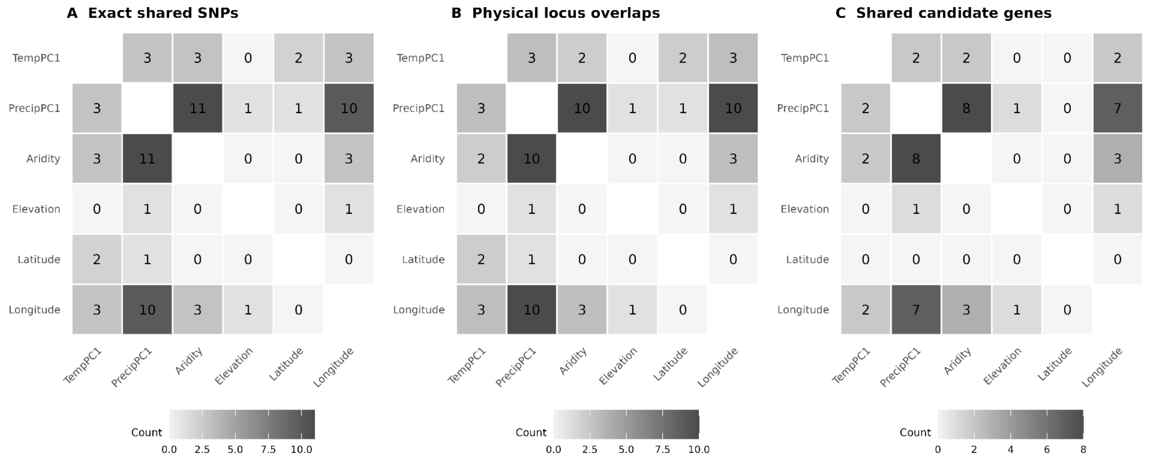


**Figure S9. Pairwise overlap among genotype–environment association candidates across environmental and spatial covariates.** Pairwise overlap among TemperaturePC1, PrecipitationPC1, aridity index, elevation, latitude, and longitude is summarized at three levels: (A) number of exact SNPs exceeding BF > 20 dB for both covariates, (B) number of physically overlapping GEA candidate-locus pairs, and (C) number of shared candidate genes. Diagonal cells are omitted. Sharing was strongest among PrecipitationPC1, aridity index, and longitude, whereas several covariate pairs showed little or no overlap.

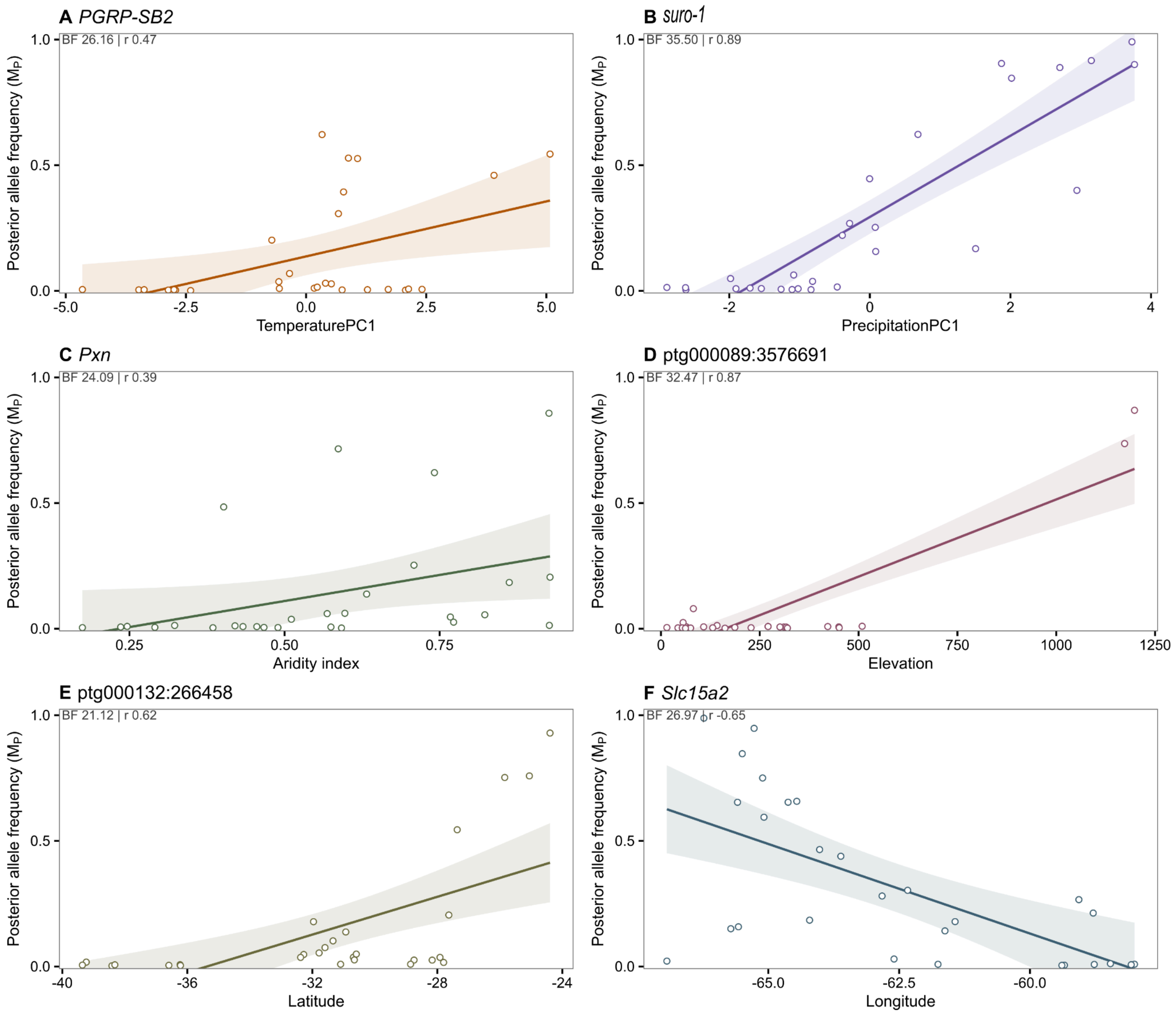


**Figure S10. Representative genotype–environment associations at non-redundant GEA-only loci.** BayPass posterior population allele frequencies ($M_P$) from the representative seed 13579 run are plotted against the corresponding environmental or spatial covariate for one representative locus per covariate: (A) *PGRP-SB2*–TemperaturePC1, (B) *suro-1*–PrecipitationPC1, (C) *Pxn*–aridity index, (D) ptg000089:3576691–elevation, (E) ptg000132:266458–latitude, and (F) *Slc15a2*–longitude. Representative loci were selected as the strongest gene-associated GEA candidates within 5 kb that did not physically overlap *XtX* or $C_2$ candidate regions and were not already displayed in the main figure. To avoid redundant representation of the same genomic region across covariates, when the strongest eligible locus was shared by more than one covariate, it was retained for the covariate with the highest median Bayes Factor and the next-highest eligible non-overlapping locus was selected for the remaining covariate(s). Candidate selection was independent of whether a curated biological gene symbol was available; biological gene symbols are shown where available, whereas genomic coordinates are shown for loci associated only with internal annotation IDs. Lines indicate linear trends with 95% confidence intervals, and inset values report the median Bayes Factor across three independent BayPass runs, expressed in decibans [BF(dB)], and the descriptive Pearson correlation coefficient (*r*). These allele-frequency patterns are intended as descriptive visualizations of the BayPass associations rather than independent tests of selection.
